# PhageLysClass: Evidence-aware classification of phage lytic proteins using protein language models

**DOI:** 10.64898/2026.11.19.767946

**Authors:** David Medina-Ortiz, Álvaro Olivera-Nappa, María Elena Lienqueo, Alejandro Villasante, Jaime Romero

## Abstract

Phage lytic proteins are promising antibacterial and biotechnological candidates, yet un-certain annotations, modular architectures, and class imbalance complicate sequence-based classification. We developed PhageLysClass, an evidence-aware framework for recognizing lytic proteins and distinguishing Endolysins, Depolymerases, and virion-associated lytic enzymes. From 442,329 unique phage sequences, including 11,867 evidence-supported positives, we defined distinct candidate non-target, unlabeled, and challenge populations. Seven supervised and two positive–unlabeled tasks were evaluated across factorial redundancy and train–test homology conditions, yielding 2,240 and 768 experiments, respectively. Redundancy reduction influenced generalization more strongly than homology separation, with non-additive interactions between them. ESM2-T30 consistently outperformed alternative representations and supported a hierarchical classifier achieving approximately 0.89 MCC under strict R30/S30 conditions in the selected, pre-refitting benchmark. Applied without retraining to 77,897 sequences across three collections, PhageLysClass combined predictions with physicochemical and geometric contextualization, highlighting Depolymerase-like candidates among structurally ambiguous proteins while distinguishing reference-supported assignments from extrapolation.

## 1 Introduction

Bacteriophages encode diverse proteins that disrupt or remodel bacterial surface barriers during infection. Endolysins hydrolyze peptidoglycan to promote host-cell lysis and progeny release, whereas virion-associated lytic enzymes (VALs) support localized cell-wall degradation during phage entry (Latka et al., 2017; Alrafaie and Stafford, 2023). Phage depolymerases instead target capsules, extracellular polysaccharides, lipopolysac-charides, and other surface glycans (Knecht et al., 2020; Cheetham et al., 2024), contributing to host recognition and biofilm disruption (Topka-Bielecka et al., 2021). These activities have made phage-derived enzymes attractive as antibacterial and biotechnological agents (Fernandes and São-José, 2018; Grishin et al., 2020; Lee et al., 2023). Their sequence-based identification remains difficult because related functions can occur in proteins with markedly different lengths, domain architectures, and global sequence similarity (Bałdysz et al., 2024; Latka et al., 2017; Alrafaie and Stafford, 2023).

Computational recognition of phage lytic and surface-degrading proteins has progressed from amino-acid composition and physicochemical descriptors (Chen et al., 2016; Meng et al., 2020) to multifeature machine learning and pretrained protein language models (Li et al., 2020). Targeted classifiers have been developed for phage depolymerase identification (Magill and Skvortsov, 2023; Concha-Eloko et al., 2024; Shen et al., 2026), while broader resources such as PhaLP and PhaLP 2.0 support the organization and prediction of phage lytic proteins (Criel et al., 2021; Boulay et al., 2026). Related studies have extended sequence-based discovery to poorly characterized lysins and large-scale depolymerase mining (Fu et al., 2024; Zhang et al., 2024; Vieira et al., 2025). Protein language models are particularly attractive because their representations capture transferable information related to protein sequence, structure, and function (Elnaggar et al., 2021; Lin et al., 2022; Elnaggar et al., 2023). However, their predictive value remains dependent on how biological targets, reference backgrounds, and evaluation datasets are defined.

This dependence is relevant for phage proteins. Endolysins, VALs, depolymerases, tail-associated proteins, and other lysis-related proteins share partially overlapping functional and architectural features, while non-target examples may range from proteins with well-supported alternative functions to sequences with unre-solved annotation (Nasiri Shoeibi et al., 2026; Grigson et al., 2025). Database coverage is also taxonomically and experimentally biased, and functional annotations vary in depth and provenance (Cook et al., 2021; Bateman et al., 2025). The absence of a lytic annotation therefore does not provide direct evidence of non-lytic function (Zhapa-Camacho et al., 2024; Grigson et al., 2025). Sequence similarity introduces a second source of uncertainty. Redundancy can inflate the apparent amount of independent training evidence, while close train–test homologs can produce optimistic estimates of generalization (Soto-García et al., 2026). Redundancy control and homology separation address different aspects of this problem but are often reduced to a single sequence-identity criterion (Soto-García et al., 2026). Their combined effects, together with biological task definition and representation choice, remain poorly characterized. Positive–unlabeled learning provides an additional formulation for the large fraction of phage sequence space whose function remains unresolved (Zhapa-Camacho et al., 2024).

Here, we present PhageLysClass, an evidence-aware framework for sequence-based recognition and functional classification of phage lytic and surface-degrading proteins. Starting from 442,329 unique phage proteins, including 11,867 evidence-supported positives from the PhageLysData Core (Medina Ortiz et al., 2026), we define positive, negative, unlabeled, and challenge states and characterize their physicochemical and representation-space organization. We evaluate seven supervised and two positive–unlabeled tasks across independently controlled redundancy and train–test homology regimes, comprising 2,240 supervised and 768 positive–unlabeled experiment families. This analysis identified ESM2-T30 as the most transferable representation and supported a compact hierarchical architecture that retained an MCC of approximately 0.89 for global lytic recognition and approximately 0.91 for Endolysin–Depolymerase–VAL multiclass classification under the strict R30/S30 condition, combining 30% sequence-identity redundancy control with 30% train–test homology separation. The frozen system was then applied without retraining to 77,897 proteins from three application collections excluded from model development, providing a controlled demonstration of lytic-protein recognition, functional classification, and contextualization against evidence-supported reference sequences.

## 2 Methods

The PhageLysClass workflow was organized into five connected stages spanning evidence definition, protein characterization and sequence-space control, matched model development, generalization analysis, and construction and application of the final predictive architecture (Figure 1). Biological evidence states and prediction tasks were established before representation generation and model fitting, while sequence redundancy and train–test homology were controlled independently throughout model development. Model selection and architecture definition were completed before any application dataset was analyzed. This separation allowed the effects of biological evidence, representation choice, learner selection, and evaluation conditions to be examined independently within a single controlled framework.

### 2.1 Evidence universe and task formulation

The positive reference universe was derived from the evidence-supported PhageLysData Core (Medina Ortiz et al., 2026), while a complementary phage-protein background was obtained from UniProtKB release 2026_02 (Bateman et al., 2025) using taxonomy-based retrieval of Caudoviricetes and named bacteriophage proteins outside this class.

**Figure 1.**
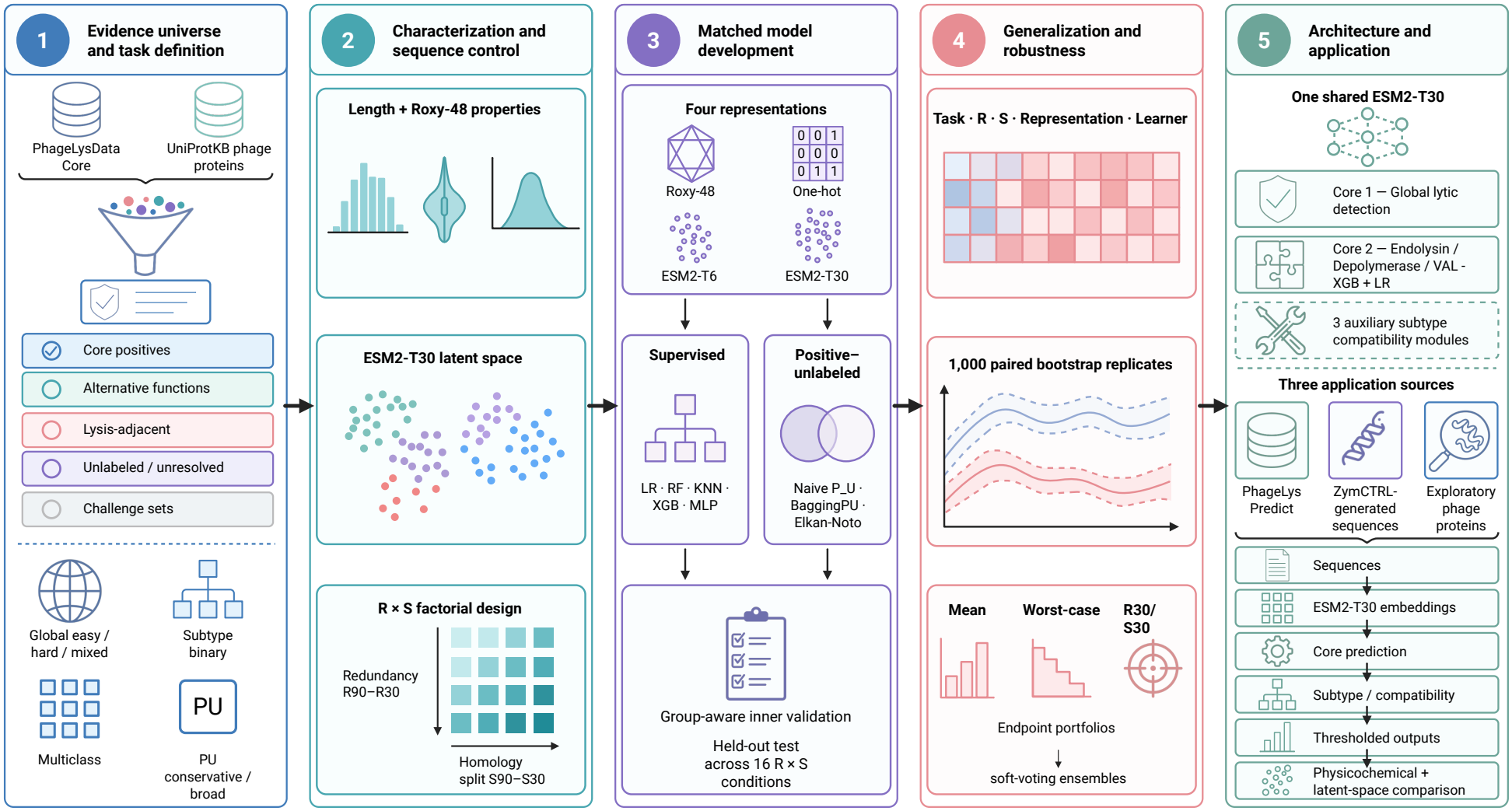
Evidence-aware development and application workflow of PhageLysClass. The frame-work begins with the construction of an evidence universe from the PhageLysData Core and UniProtKB phage proteins. Positive, alternative-function, lysis-adjacent, unlabeled, and challenge sequence states are then defined and converted into global, subtype-specific, multiclass, and positive–unlabeled prediction tasks. Protein properties and ESM2-based representation spaces are characterized before model development. Sequence redundancy (*R*) and train–test homology (*S*) are controlled independently through a factorial design. Four protein representations and multiple supervised and positive–unlabeled learning strategies are compared under matched sequence assignments. Generalization is then evaluated across the complete task *× R × S ×* representation *×* learner landscape. The selected architecture uses a shared ESM2-T30 representation for global lytic recognition, functional classification, and auxiliary subtype compatibility. After model selection and decision rules are frozen, the system is applied without retraining to the PhageLys Prediction Extension, ZymCTRL-generated proteins, and an ambiguous structural challenge set. Predictions are then contextualized against evidence-supported reference proteins using physicochemical properties and reference-space geometry.

UniProtKB proteins were assigned to evidence states using a frozen hierarchy based on protein names, catalytic annotations, enzyme classifications, keywords, and curated functional comments. Exact sequence matches to the PhageLysData Core were retained as known positive overlaps. Proteins with explicit alternative functions formed the principal non-lytic background, whereas holins, antiholins, spanins, and related proteins formed a separate lysis-adjacent state. Proteins lacking sufficient functional support remained unlabeled, while structurally or functionally ambiguous proteins were retained as challenge populations.

The frozen evidence universe was then converted into complementary prediction tasks before redundancy reduction, representation generation, or model fitting. Three global binary tasks compared the complete positive Core with the alternative-function background, the lysis-adjacent background, or their union. Independent binary tasks were defined for Endolysin, Depolymerase, and virion-associated lytic enzyme recognition using the combined explicit non-lytic background, while a clean multiclass task contained only single-class Endolysin, Depolymerase, and VAL sequences. Two positive–unlabeled formulations were also constructed from known positives and unresolved proteins. Unlabeled sequences were retained as unknown rather than converted into biological negatives, and challenge populations remained outside ordinary model fitting. Supplementary Information, Section S1.1 provides the complete source-integration procedures, evidence-state assignment hierarchy, annotation audit, and detailed composition of the nine prediction tasks.

### 2.2 Protein characterization and representation-space analysis

The frozen sequence universe was characterized before predictive modelling using sequence length, amino-acid composition, and physicochemical properties calculated with Roxy (Medina-Ortiz et al., 2026a). Initial characterization included 71 composition-related descriptors and 27 physicochemical descriptors, analysed separately and jointly in a 98-descriptor representation. The subsequent post-prediction contextualization used 26 physicochemical descriptors after excluding molecular weight, whereas predictive benchmarking em-ployed a distinct 48-feature Roxy representation. Core positives were compared with the alternative-function, lysis-adjacent, and combined supervised backgrounds, while clean Endolysin, Depolymerase, and VAL sequences were analysed separately. Group differences were quantified using Cliff’s delta and non-parametric tests with Benjamini–Hochberg correction, emphasizing effect size and consistency across biological back-grounds.

Multivariate organization was evaluated in explicit descriptor spaces and pretrained protein-language-model spaces using ESM2-T6 and ESM2-T30 representations (Lin et al., 2022). PCA, UMAP, and t-SNE were used for visualization, while quantitative analyses were performed in the corresponding original descriptor or embedding spaces using global separation and local-neighborhood measures. These analyses did not modify task membership or model selection. Supplementary Information, Section S1.2 provides the complete descriptor selection procedures, characterization spaces, statistical analysis configurations, balanced sampling strategies, and parameters used for geometric analysis and dimensionality reduction.

### 2.3 Sequence redundancy and homology-aware evaluation

Sequence redundancy was controlled before train–test partitioning. For each prediction task, four independently reduced datasets were generated at sequence-identity thresholds denoted R90, R70, R50, and R30. Reduction was performed separately within the biological groups defining each task before retained sequences were recombined, preserving the roles of positive, negative, and unlabeled populations.

Each redundancy-controlled dataset was subsequently subjected to homology-aware partitioning at four sequence-identity thresholds denoted S90, S70, S50, and S30. Thus, every redundancy regime was evaluated under four distinct train–test homology conditions. Sequence groups were generated with MMseqs2 (Steinegger and Söding, 2017) and assigned to development or held-out test partitions using BioSieve homology-aware splitting (Villegas et al., 2026). Whole sequence groups were kept intact across the partition boundary, with approximately 30% of each redundancy-controlled dataset retained for outer testing.

This sequential procedure produced 16 *R × S* conditions for each prediction task, with R90, R70, R50, and R30 each evaluated under S90, S70, S50, and S30. The design therefore quantified how train–test homology affected generalization within progressively stricter redundancy-controlled sequence spaces. Representation and learner comparisons within each condition used the same stored protein assignments. Supplementary Information, Section S1.3 documents the redundancy-reduction and homology-clustering configurations, cluster-preserving partition procedures, assignment corrections, and train–test integrity audits.

### 2.4 Matched model development

Four protein representations were compared under matched task and sequence-space assignments. These comprised 48 Roxy sequence-derived descriptors, a fixed-length one-hot encoding, and mean-pooled ESM2-T6 and ESM2-T30 embeddings generated with Sylphy (Medina-Ortiz et al., 2026a; Lin et al., 2022; Medina-Ortiz et al., 2026b).

Five supervised learning algorithms were evaluated for each representation, including logistic regression, random forest, *k*-nearest neighbors, XGBoost, and multilayer perceptron models. Candidate configurations were selected using group-aware inner validation restricted to the development partition. Binary model selection prioritized MCC followed by average precision, while multiclass selection prioritized multiclass MCC followed by macro-averaged F1 score. Selected configurations were refitted on the complete development partition and evaluated once on the corresponding held-out test set.

The supervised benchmark varied biological task, redundancy regime, homology regime, representation, and learner while preserving matched sequence assignments for direct comparisons. Predictions, probability estimates, model metadata, and test membership were retained for downstream robustness analyses.

Positive–unlabeled learning was evaluated separately using naive positive-versus-unlabeled learning, BaggingPU, and Elkan–Noto correction with controlled base learners (Nanduri et al., 2026). Evaluation considered both recovery of held-out known positives and ranking behaviour within the unlabeled population. Supplementary Information, Section S1.4 provides the representation eligibility and preprocessing criteria, complete learner configurations, group-aware validation procedures, and detailed training and evaluation protocols for the positive–unlabeled strategies.

### 2.5 Generalization, robustness, and architecture selection

Generalization was examined through individual redundancy and homology contrasts, their interactions across the *R × S* landscape, and common-test comparisons restricted to proteins shared between evaluation conditions. Principal matched contrasts were evaluated using 1,000 paired bootstrap replicates to quantify uncertainty and distinguish changes in prediction behaviour from differences in test-set composition.

Robustness was assessed across all 16 *R × S* conditions. Each task–representation–learner strategy was summarized using mean performance, worst-case performance, strict R30/S30 performance, performance range, and ranking stability. Strategies were shortlisted only after completion of the full benchmark, considering predictive performance, robustness, uncertainty, and computational cost without combining these criteria into a single score.

For each prediction endpoint, retained strategies were evaluated individually and in compatible probability-level ensembles across the same sequence-space conditions. Unweighted soft voting was used as the primary ensemble formulation, and ensembles were retained only when they provided a reproducible improvement over their constituent models. Final model composition was determined independently for global lytic recognition, multiclass functional classification, and the subtype-specific auxiliary components.

After architecture selection, representation choice, model membership, voting rules, and decision procedures were frozen. Final production models were refitted using all eligible data from their corresponding task definitions, and application datasets were not used to revise any selection or decision rule. Supplementary Information, Section S1.5 provides the mathematical definitions of the performance and robustness measures, common-test comparisons, bootstrap uncertainty estimation, candidate shortlisting criteria, ensemble construction and selection procedures, and final production-model configuration.

### 2.6 Application of the frozen PhageLysClass system to heterogeneous sequence collections

The frozen PhageLysClass architecture was applied to three heterogeneous sequence collections excluded from model development. These comprised the PhageLys Prediction Extension (Medina Ortiz et al., 2026), de novo proteins generated with ZymCTRL (Munsamy et al., 2022) under enzyme-specific conditioning, and UniProtKB phage proteins with ambiguous structural annotations. These collections provided complementary application scenarios involving computationally annotated, artificially generated, and functionally ambiguous proteins.

All sequences were processed using the frozen ESM2-T30 representation and hierarchical predictive architecture. Global lytic recognition preceded conditional Endolysin, Depolymerase, and VAL classification, while auxiliary subtype models provided independent compatibility outputs. Supplementary Information, Section S1.6 details the preparation of each collection, ZymCTRL generation settings, sequence-quality controls, and frozen inference procedures.

Predictions were subsequently contextualized against evidence-supported reference proteins using 26 sequence-derived physicochemical descriptors. Compatibility with global and subtype-specific reference populations was assessed through descriptor distributions, empirical distance boundaries, and local neighborhood organization in the complete standardized descriptor space. These complementary criteria were integrated to distinguish concordant, mixed, and discordant predictions without modifying their functional assignments. Dimensionality reduction was used exclusively for visualization. Supplementary Information, Section S1.7 provides the mathematical definitions, reference-space calculations, support thresholds, and concordance criteria.

## 3 Results and Discussion

### 3.1 Biological and representational organization of the PhageLysClass evidence universe

Before model development, we examined how the evidence-supported lytic proteins were positioned within the broader phage-protein sequence universe. The resulting landscape was highly heterogeneous, with marked differences in evidence support, sequence architecture, composition, and physicochemical properties across biological groups. These differences were not equally preserved across comparison backgrounds, and their organization became clearer when sequences were represented with pretrained protein language models. This initial characterization therefore provided the biological context needed to interpret the predictive tasks and the generalization behavior evaluated later.

#### 3.1.1 Evidence-aware construction reveals a predominantly unresolved phage-protein universe

Integration of the evidence-supported PhageLysData Core with a taxonomy-defined UniProtKB phage-protein background yielded 442,329 unique protein sequences. UniProtKB retrieval produced 570,195 accession-level records, corresponding to 437,447 unique sequences after exact-sequence deduplication. Of the 11,867 PhageLysData Core sequences, 6,985 were also found in UniProtKB, while 4,882 were exclusive to PhageLysData. The final universe therefore represents the union of both collections, with shared sequences counted only once.

The resulting universe was dominated by proteins with insufficient functional evidence to establish their relationship to phage lytic activity (Figure 2A). Each sequence was assigned to a mutually exclusive primary evidence state. All percentages describing universe-level composition were calculated relative to the 442,329 unique sequences. The evidence-supported positive Core comprised 11,867 sequences (2.7%), while candidate non-target proteins accounted for 59,038 sequences (13.3%). The latter included 54,711 proteins with supported alternative functions (easy negatives) and 4,327 proteins associated with lysis-adjacent mechanisms (hard negatives). These two populations were maintained separately and combined only for the mixed supervised background. Neither was assumed to represent experimentally verified absence of lytic activity.

The remaining 371,424 sequences (84.0%) comprised 360,220 unlabeled or otherwise unresolved proteins and 11,204 proteins assigned to lytic-related or structurally ambiguous challenge populations. Unlabeled proteins lacked sufficient evidence for positive or negative assignment, whereas challenge proteins carried annotations that prevented their unambiguous classification within the principal evidence states. The predominance of these populations highlights the limited functional characterization of the broader phage-protein universe. It also supports their separation from candidate negatives during predictive model development.

The positive Core exhibited substantial functional class imbalance. Of its 11,867 sequences, 10,841 had a single assignment to one of the three principal lytic classes. This clean single-label subset comprised 9,888 Endolysins (91.2%), 712 Depolymerases (6.6%), and 241 virion-associated lytic enzymes (VALs; 2.2%). These percentages were calculated relative to the 10,841 single-label sequences. The remaining 1,026 Core proteins included 999 with multiple non-exclusive lytic assignments and 27 associated with other phage envelope lytic functions. Thus, the principal subtype classification problem was dominated by Endolysins, with substantially fewer examples supporting Depolymerase and VAL recognition.

Protein length further distinguished the evidence-defined populations (Figure 2B). The complete positive Core had a median length of 290 residues, compared with 334 residues for the alternative-function background and 112 residues for the lysis-adjacent background. Within the clean single-label Core, median lengths were 278 residues for Endolysins, 375 for VALs, and 692 for Depolymerases. Approximately 10.4% of Depolymerases exceeded 1,024 residues, compared with 5.2% of Endolysins and less than 1% of VALs. These proportions were calculated within each subtype. T hese differences indicate that the prediction tasks varied not only in their supporting biological evidence but also in their underlying sequence architecture.

#### 3.1.2 Lytic proteins show reproducible sequence-property signatures with strong background dependence

Core proteins differed systematically from the supervised backgrounds in amino-acid composition (Figure 2C). Glycine showed the strongest background-stable enrichment, with Cliff’s delta values of 0.62 against the alternative-function background, 0.55 against the lysis-adjacent background, and 0.62 against the mixed background. Tryptophan was also enriched across the three comparisons. Leucine showed the opposite pattern and remained depleted, with its strongest effect against the lysis-adjacent population at *δ* = *−*0.61.

Several compositional effects depended strongly on the comparison background. Glutamate was markedly depleted relative to alternative-function and mixed proteins, with *δ* = *−* 0.62 and *−* 0.57, but showed little difference from lysis-adjacent proteins. Histidine, proline, aspartate, and arginine became more discriminative against the lysis-adjacent background. Aspartate changed direction between comparisons and was depleted relative to alternative-function proteins but enriched relative to lysis-adjacent proteins. The apparent compositional signature of lytic function therefore changed with the biological definition of the non-target population.

**Figure 2.**
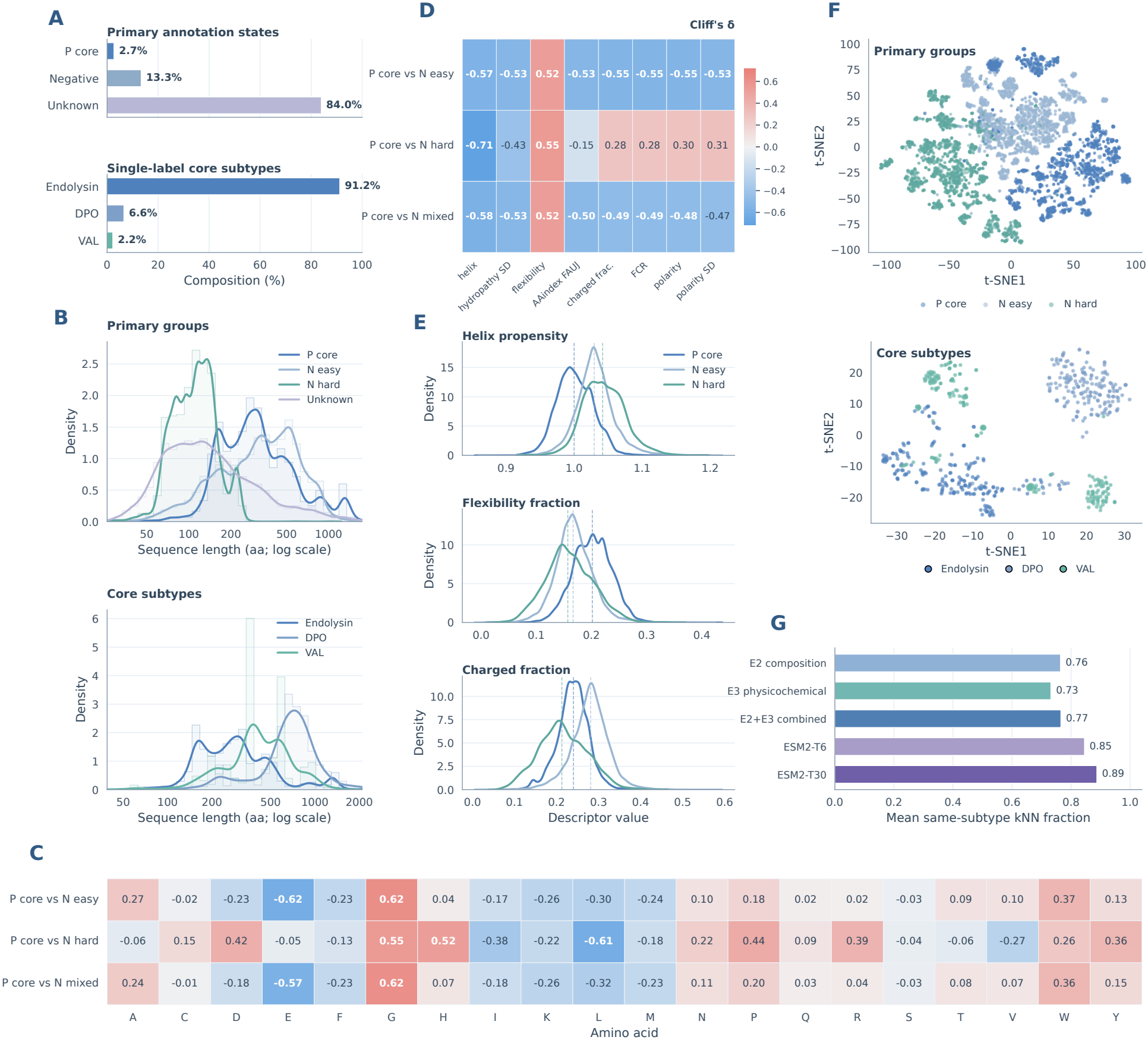
Sequence-level and representation-space organization of the PhageLysClass evidence universe. **A**, composition of the primary evidence states and of the clean single-label PhageLysData Core used for subtype analysis. **B**, sequence-length distributions for the primary evidence groups and the three major lytic subtypes. **C**, amino-acid composition differences between Core positives and the alternative-function, lysis-adjacent, and mixed negative backgrounds, expressed as Cliff’s delta. **D**, physicochemical differences between Core positives and the three supervised backgrounds, expressed as Cliff’s delta. **E**, distributions of selected physicochemical descriptors illustrating stable and background-dependent effects. **F**, t-SNE projections of ESM2-T30 representations showing the organization of primary evidence states and clean Endolysin, Depolymerase, and VAL populations. **G**, local functional organization across five representation spaces: E2 (amino-acid sequence composition descriptors), E3 (physicochemical descriptors), E2+E3 (combined sequence composition and physicochemical descriptors), ESM2-T6, and ESM2-T30 (pretrained protein language model embeddings). Local subtype consistency was quantified as the mean fraction of nearest neighbors sharing the same lytic subtype at *k* = 25. Dataset identifiers N easy, N hard, and N mixed correspond to the alternative-function, lysis-adjacent, and combined negative backgrounds, respectively.

Physicochemical properties showed a similar combination of stable and background-dependent effects (Figure 2D,E). Helix propensity was consistently lower in Core proteins, with Cliff’s delta values between *−* 0.57 and *−* 0.71. Flexibility remained higher across all backgrounds, with effect sizes between 0.52 and 0.55. These properties therefore distinguished Core proteins under substantially different negative definitions.

Charge-related properties were less transferable across backgrounds. Charged fraction was lower in Core proteins than in the alternative-function population, with *δ* = *−* 0.55, but higher than in the lysis-adjacent population, with *δ* = 0.28. Related polarity and electrostatic descriptors showed comparable changes in direction or magnitude. The mixed population largely followed the alternative-function profile because alternative-function proteins represented most of the explicit negative universe.

These results do not support a single physicochemical definition of phage lytic proteins. Some sequence properties were stable across biological backgrounds, while others were specific to the comparison population. Such background dependence is important for predictive modelling because strong discrimination within one dataset may reflect properties of the selected negative class without necessarily indicating a transferable signal of lytic function.

#### 3.1.3 Protein language model representations strengthen functional organization without creating discrete sequence classes

The ESM2-T30 t-SNE projections revealed preferential organization of evidence states and functional sub-types without forming completely separated populations (Figure **2**F). Within the positive Core, Depoly-merases and VALs occupied more localized regions, whereas Endolysins showed a broader distribution. Nevertheless, substantial overlap and transitional regions remained, indicating that functional annotations did not correspond to discrete boundaries in the projected representation space.

Local neighborhood analysis provided quantitative support for this organization (Figure **2**G). The mean fraction of same-subtype nearest neighbors at *k* = 25 was 0.76 for amino-acid composition, 0.73 for physic-ochemical descriptors, and 0.77 for their combined representation. This fraction increased to 0.85 with ESM2-T6 and 0.89 with ESM2-T30. Within ESM2-T30 space, local subtype coherence was highest for Depolymerases, followed by VALs and Endolysins. These results indicate that protein language model representations captured functional relationships more consistently than explicit sequence descriptors, although subtype organization remained heterogeneous.

Complementary silhouette analyses assessed the global separation of Core positives from the supervised non-target backgrounds. Against the alternative-function background, silhouette scores were 0.048 for amino-acid composition, 0.082 for physicochemical descriptors, and 0.058 for their combination. These values increased to 0.101 with ESM2-T6 and 0.144 with ESM2-T30. A similar pattern was observed against the mixed background, with silhouette scores of 0.097 and 0.139 for ESM2-T6 and ESM2-T30, respectively. Thus, stronger local functional organization in protein language model spaces was accompanied by improved, but still limited, global separation between lytic and non-target proteins.

An unexpected pattern emerged for the lysis-adjacent background. Despite its biological proximity to the targeted lytic functions, this population showed greater separation from Core positives than either the alternative-function or mixed backgrounds across the evaluated representations. With ESM2-T30, the sil-houette score reached 0.312, compared with 0.144 and 0.139 for the alternative-function and mixed back-grounds, respectively. Biological proximity therefore did not necessarily imply greater similarity in numerical representation spaces. These findings also indicate that biological task definitions alone cannot establish representational difficulty or predict downstream classification performance. Further results on global separation across representations and non-target backgrounds are provided in Supplementary Results Section S2.1.

### 3.2 Sequence-space control and predictive generalization define the PhageLysClass architecture

The predictive benchmark was designed to determine which aspects of model performance were stable across changes in sequence redundancy, train–test relatedness, representation, and learner choice. Generalization varied substantially across this landscape, revealing that apparent model strength depended not only on the biological task but also on how sequence space was controlled. These comparisons ultimately favored representations and model combinations that remained competitive under increasingly restrictive conditions and provided the basis for the frozen PhageLysClass architecture.

#### 3.2.1 Generalization depends strongly on task definition and sequence-space control

The supervised benchmark comprised 2,240 completed experiment families spanning seven prediction tasks, four redundancy regimes, four train–test homology regimes, four protein representations, and five down-stream learners. Performance varied substantially among biological tasks (Figure **3**A). Global recognition against the lysis-adjacent background showed the strongest performance. Depolymerase recognition and clean multiclass classification also occupied high-performance regimes. Global recognition against alternative-function and mixed backgrounds and Endolysin recognition showed intermediate performance. VAL was the most variable endpoint. Extended task-level performance analyses are reported in Supplementary Results Section S2.2.1.

The lysis-adjacent task confirmed the pattern observed during representation-space characterization. Across matched conditions, its median MCC exceeded the alternative-function task by approximately 0.17. The mixed and alternative-function formulations differed by only approximately *−* 0.004. Analyses restricted to proteins shared between compared test sets produced the same qualitative result. The difference was therefore not explained only by changes in test-set composition. The mixed formulation was retained as the principal global task because it represented the complete explicit non-target population.

Redundancy reduction had a larger effect on observed predictive performance than train–test homology separation (Figure **3**C). In native-test comparisons, moving from R90 to R30 produced a median MCC change of approximately *−* 0.075 across the supervised benchmark. The corresponding S90-to-S30 change was approximately *−* 0.02. The redundancy effect varied by task. Global alternative-function and mixed recognition lost approximately 0.10 MCC, while Endolysin recognition decreased by approximately 0.14. The lysis-adjacent task lost approximately 0.05. Depolymerase recognition remained comparatively stable and showed a small apparent improvement. Redundancy-dependent evidence retention and sequence-space changes are reported in Supplementary Results Section S2.2.2.

The observed performance reduction was substantially smaller than the reduction in available sequences. At R30, the median retained fraction of training entities was approximately 30.7%, while 56.0% of the original homology-cluster richness was preserved. Redundancy reduction therefore decreased sequence counts more rapidly than the diversity of represented homology groups. However, comparisons between redundancy regimes also involved changes in test-set composition. When models were evaluated on shared test proteins, the median R90-to-R30 MCC difference decreased from approximately *−* 0.075 to *−* 0.012. This attenuation indicates that differences in evaluation-population composition contributed substantially to the apparent redundancy penalty, although smaller performance differences persisted for some tasks. Notably, the small native-test improvement observed for Depolymerase recognition became slightly negative under common-test evaluation.

Homology separation produced a smaller and more consistent performance penalty, although its magnitude depended on the redundancy regime. At R90, moving from S90 to S30 reduced MCC by approximately 0.047 in native-test comparisons and 0.037 on shared test proteins. At R30, both effects approached zero. Thus, the influence of train–test homology separation diminished as redundancy control became more stringent, and this attenuation persisted under common-test evaluation. The corresponding native-test and common-test comparisons are detailed in Supplementary Results Section S2.2.3.

The *R × S* response surfaces further showed that the two sequence-control factors were not additive. Across the extreme R90/R30 and S90/S30 contrast, the median interaction was approximately +0.051 MCC, with approximately 89% of task–representation–learner surfaces showing a positive interaction. Because both main effects generally reduced performance, the positive interaction indicates that their combined effect was less detrimental than expected from their isolated effects. This pattern is consistent with partially overlapping constraints on sequence-space generalization. Importantly, these interaction estimates describe the original benchmark surfaces, whose test populations differed across conditions. The limited number of proteins shared across all four extreme conditions prevented a reliable equivalent interaction analysis on identical test sequences. The interaction should therefore be interpreted as a property of the factorial evaluation landscape, while common-test comparisons provide more direct evidence of prediction differences on shared proteins. The complete interaction analysis is reported in Supplementary Results Section S2.2.3.

**Figure 3.**
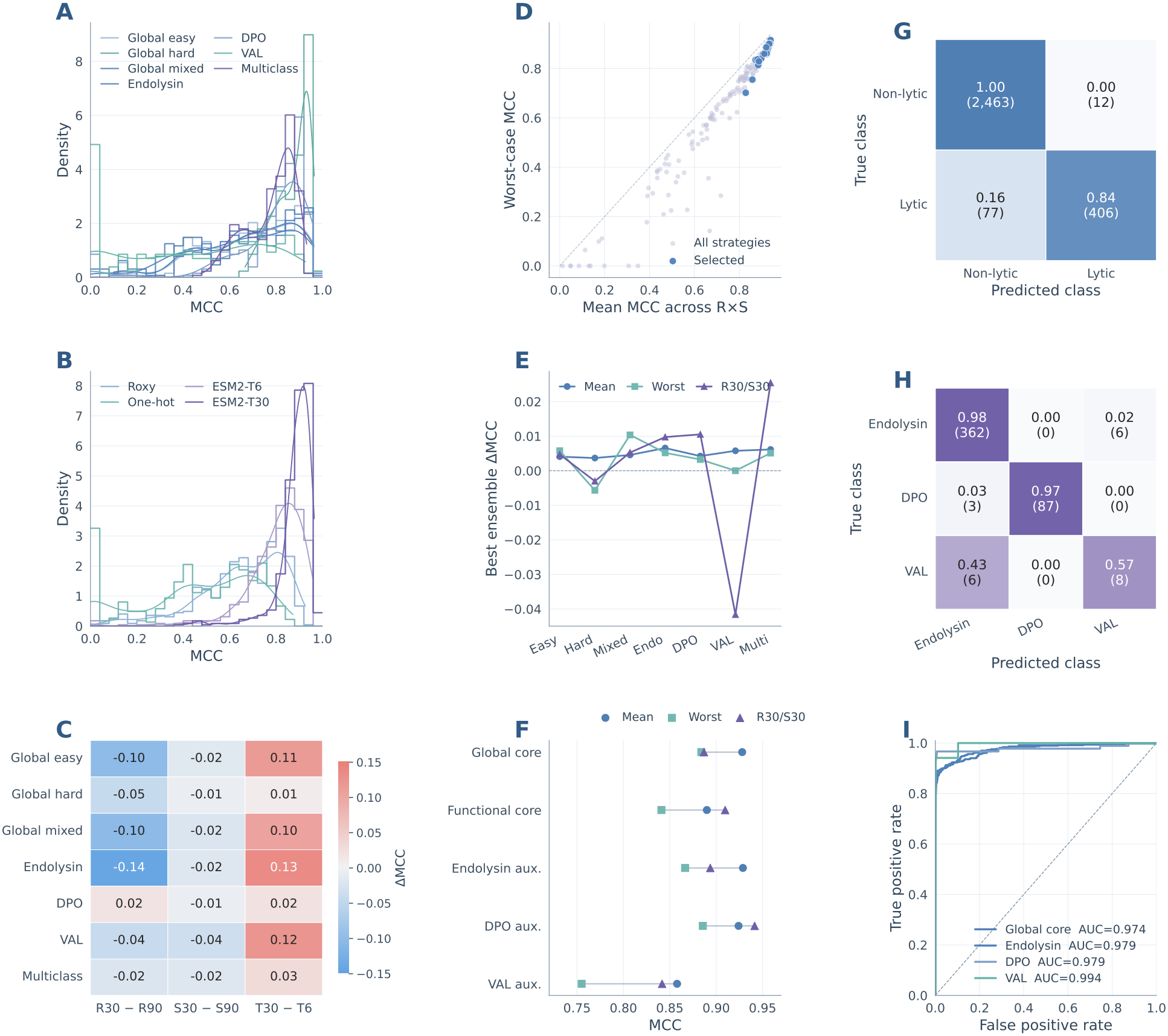
Matched benchmarking and derivation of the PhageLysClass predictive architecture. **A**, performance regimes across the supervised prediction tasks. **B**, comparative performance of the four protein representations across matched experimental conditions. **C**, effects of redundancy reduction, train–test homology separation, and increased ESM2 representation capacity on predictive performance. **D**, robustness-based model-selection space relating mean and worst-case performance across the complete *R × S* landscape. **E**, endpoint-specific performance changes obtained by combining retained models through soft-voting ensembles. **F**, performance of the final core and auxiliary PhageLysClass components across sequence-space conditions. **G**, confusion matrix for the final global lytic-recognition component under the strict R30/S30 condition. **H**, confusion matrix for the final Endolysin–Depolymerase–VAL multiclass component under the strict R30/S30 condition. **I**, ROC curves for the final binary components under strict sequence-space control.

#### 3.2.2 Representation capacity is more transferable than downstream learner choice

Protein representation was one of the most consistent determinants of generalization (Figure **3**B,C). ESM2-T30 produced the strongest overall performance distribution and showed the greatest stability across task definitions and sequence-space conditions. Its advantage over ESM2-T6 was largest for tasks that remained representation limited. Median MCC increased by approximately 0.11 for global alternative-function recognition, 0.10 for global mixed recognition, 0.13 for Endolysin, and 0.12 for VAL. The improvement was smaller for the already highly separable lysis-adjacent and Depolymerase tasks.

Robustness showed the same representation hierarchy. Sensitivity to biological task definition and sequence-space restriction generally decreased from one-hot encoding to Roxy, ESM2-T6, and ESM2-T30. The dependence of apparent task difficulty on the negative background also decreased as representation capacity increased. In prevalence-normalized average-precision skill, the median advantage of the lysis-adjacent task over the alternative-function task decreased from approximately +0.48 with one-hot encoding to +0.24 with Roxy, +0.09 with ESM2-T6, and +0.02 with ESM2-T30. The stronger protein language model therefore reduced the extent to which apparent learnability depended on the chosen non-target population. Extended representation analyses are reported in Supplementary Results Section S2.3.1.

Downstream learner effects were less transferable. No algorithm dominated across prediction tasks and sequence-space conditions. Learner rankings changed with endpoint, representation, and evaluation regime. Differences among competitive learners were generally smaller than those produced by representation choice. Learner-specific robustness and ranking stability are reported in Supplementary Results Section S2.3.2.

Positive–unlabeled learning showed a related but distinct pattern. Across 768 PU experiment families, stronger protein representations supported comparable recovery of known positives under naive positive-versus-unlabeled learning, BaggingPU, and Elkan–Noto correction. Larger differences appeared in the ordering of unresolved sequences. Elkan–Noto correction largely preserved the ranking produced by naive learning while changing score scaling. BaggingPU could produce substantially different rankings. Recovery of known positives and prioritization within unresolved sequence space therefore represented different operating objectives. PU learning was retained as a separate discovery-oriented analysis and was not in-corporated into the supervised PhageLysClass core. Extended PU analyses are reported in Supplementary Results Section S2.3.3.

#### 3.2.3 Endpoint-specific model combination produces a compact hierarchical architecture

Robustness analysis across 140 task–representation–learner profiles identified 23 competitive predictive strategies (Figure **3**D). Of these, 20 used ESM2-T30 and three used ESM2-T6. Neither one-hot encoding nor Roxy descriptors appeared among the retained strategies. Although ESM2-T30 was consistently favored, the strongest learner combinations differed among prediction endpoints. This indicated that representation quality was comparatively transferable across tasks, while the benefits of individual learning algorithms depended on the functional classification problem.

Ensembling produced measurable but heterogeneous performance gains (Figure **3**E). The selected global lytic-recognition ensemble achieved a mean MCC of 0.9280 across the R–S conditions, improving the strongest individual-model baseline by 0.0045 MCC on average. Its worst-case MCC increased by 0.0104, indicating that model combination contributed more strongly to robustness than to average performance. The clean multiclass ensemble achieved a mean MCC of 0.8899, with a modest average gain of 0.0028 but a larger improvement of 0.0197 under R30/S30. This was the largest strict-condition gain among the selected ensembles.

Subtype-specific ensembles displayed different trade-offs. The Endolysin ensemble achieved a mean MCC of 0.9288, improving average, worst-case, and strict-condition performance by 0.0066, 0.0034, and 0.0048 MCC, respectively. Depolymerase recognition reached a mean MCC of 0.9241 and gained 0.0105 under R30/S30, although its worst-case performance was marginally lower than that of the strongest individual model (ΔMCC = *−* 0.0002). VAL showed the clearest limitation of model combination. Adding logistic regression to the MLP increased mean MCC by approximately 0.0053 but reduced strict-condition MCC by 0.0416. Consequently, the selected VAL component remained a single MLP, with a mean MCC of 0.8578. These results demonstrate that modest average improvements from ensembling did not necessarily translate into greater robustness under restrictive sequence-space conditions.

The resulting architecture combined two primary predictive cores with three auxiliary subtype-compatibility components. Global recognition used a KNN–MLP ensemble, while clean multiclass assignment used XG-Boost and logistic regression. The Endolysin and Depolymerase components retained their respective ensembles, whereas VAL used the standalone MLP. All five components shared ESM2-T30 embeddings, allowing the complete system to operate with one numerical representation and ten fitted estimators. The architecture therefore preserved endpoint-specific learner combinations while maintaining a common representation across the prediction hierarchy.

The selected benchmark recipes were further examined under R30/S30, the most restrictive sequence-space condition (Figure **3**F–I). Global lytic recognition correctly identified 406 of 483 Core positives (84.1%) and classified only 12 of 2,475 candidate non-target proteins as lytic (0.5%), yielding an MCC of approximately 0.89 (Figure **3**G). This result combined strong rejection of candidate non-target proteins with incomplete recovery of known positives, as 77 Core proteins remained undetected. The clean multiclass model correctly assigned 362 of 368 Endolysins (98.4%) and 87 of 90 Depolymerases (96.7%). VAL recognition was substantially less reliable, with only eight of 14 proteins (57.1%) correctly assigned and the remaining six classified as Endolysins (Figure **3**H). Thus, high classification accuracy for the dominant subtypes coexisted with considerably weaker recognition of the smallest functional class.

Binary discrimination analyses revealed an additional distinction between ranking performance and functional assignment (Figure **3**I). Under R30/S30, ROC-AUC values were approximately 0.97 for global and Endolysin recognition, 0.98 for Depolymerase, and 0.99 for VAL. The high VAL ROC-AUC was obtained from its separate binary compatibility task, whereas the eight correctly classified VAL proteins belonged to the clean multiclass evaluation. These measurements therefore describe different prediction endpoints. In particular, strong binary ranking did not eliminate the difficulty of assigning VALs within the three-class formulation. Interpretation of the VAL results was further constrained by the limited positive support, comprising 17 proteins in the strict binary test and 14 in the multiclass test.

The results supported a hierarchical architecture in which the principal recognition and functional-assignment stages were complemented by independent subtype-compatibility outputs. The observed ensemble advantages were endpoint-specific and were not uniformly preserved under strict sequence-space conditions. Importantly, the reported R30/S30 measurements correspond to the selected benchmark recipes before full-data production refitting. Because these results contributed to architecture selection, they do not constitute independent post-selection validation of the deployed models. Extended comparisons and production-model specifications are provided in Supplementary Results Section S2.4.

### 3.3 Application of the frozen architecture reveals distinct predictive and reference-space regimes

Once the predictive architecture had been frozen, we examined how it behaved across sequence collections that differed substantially from the data used during model development. The resulting predictions varied strongly by source, both in the fraction of proteins classified as lytic and in their functional composition. Comparison with physicochemical and reference-space properties further showed that some predictions remained well supported by the evidence-derived protein landscape, whereas others represented clear extrapolations beyond it.

#### 3.3.1 Prediction landscapes differ across heterogeneous sequence collections

The frozen architecture was applied to three collections excluded from model development. After source-specific processing, the analysis comprised 58,284 non-redundant proteins from the PhageLys Prediction Extension, 10,000 ZymCTRL-generated proteins, and 9,613 proteins from the ambiguous structural challenge set. These collections represented extension into a computationally annotated phage-protein universe, evaluation of generated enzyme-like sequences, and analysis of structurally related phage proteins with un-resolved functional assignments.

Global predictions differed sharply among the three sources (Figure **4**A). PhageLysClass classified 47,797 Prediction Extension proteins as lytic, corresponding to 82.0% of that collection. The positive fraction decreased to 69.5% for ZymCTRL and 44.1% for the ambiguous structural collection. These fractions describe the behaviour of the frozen model on the three application populations and do not represent performance estimates because the collections were not constructed as conventional labeled test sets.

Subtype composition also differed substantially by source (Figure **4**B). Prediction Extension positives were dominated by Endolysin assignments. Of 47,797 lytic predictions, 47,445 were assigned as Endolysins, 12 as Depolymerases, and 340 as VALs. ZymCTRL produced 6,329 Endolysin, 490 Depolymerase, and 133 VAL predictions. The ambiguous structural set showed a different profile. Depolymerases represented 2,848 of 4,235 positive predictions, compared with 1,010 Endolysins and 377 VALs.

ZymCTRL provided a controlled generative comparison. Approximately 78.9% of sequences conditioned on peptidoglycan-degrading EC classes were classified as lytic, compared with 60.2% of sequences conditioned on surface-polysaccharide-degrading classes. The generating EC and the predicted PhageLysClass subtype were not expected to correspond directly. ZymCTRL conditioning specifies catalytic activity, whereas the PhageLysClass classes describe functional roles within phage infection. The experiment therefore evaluates whether generated enzyme-like proteins enter the learned lytic decision space and does not constitute a ground-truth subtype benchmark.

The ambiguous structural collection provided a different challenge. Tailspike proteins showed the strongest enrichment for positive predictions, with approximately 90.8% classified as lytic. Most positive tailspikes were assigned as Depolymerases. Positive tail-fiber predictions also showed a strong Depolymerase tendency, while baseplate-associated proteins produced a more heterogeneous subtype distribution. This pattern is consistent with the functional diversity of virion-associated proteins involved in host-surface recognition and degradation. The assignments identify sequence groups with strong similarity to the learned Depolymerase regime but do not establish enzymatic function.

#### 3.3.2 Reference-space analysis separates supported predictions from extrapolative assignments

Physicochemical comparison revealed major differences in how closely predicted proteins resembled evidence-supported references. Prediction Extension proteins classified as lytic showed strong compatibility with the global positive population. The median sequence retained 25 of 26 frozen descriptors within the corresponding 95% reference intervals. Prediction Extension Endolysins showed a similar pattern. These descriptors were calculated independently of the ESM2-T30 representation used for classification. The agreement there-fore indicates that the dominant predictions from this source remained within the physicochemical regime represented by evidence-supported lytic proteins.

ZymCTRL sequences were less compatible with natural reference populations. Predicted lytic sequences retained a median of 16 of 26 descriptors within the global positive intervals. Predicted Depolymerases retained 13 descriptors within their subtype-specific intervals, while predicted VALs retained only five. Several of the largest deviations involved sequence-complexity measures. Generated proteins could therefore enter the model’s lytic decision space without reproducing the broader physicochemical organization of natural reference proteins.

The ambiguous structural population showed a different pattern. Proteins assigned as Depolymerases retained a median of 25 of 26 descriptors within the corresponding reference intervals. VAL assignments also showed substantial physicochemical compatibility, while Endolysin assignments were less tightly matched. This agreement is notable because the collection was defined from structural annotation and not from the PhageLysClass prediction. The ESM2-based classifier and the complementary descriptor-based analysis therefore converged on a substantial Depolymerase-like population.

Full-space reference geometry reinforced these source-dependent differences. Among Prediction Extension proteins, 59.2% of lytic predictions fell within the empirical 95% coverage of the global positive reference, compared with 19.5% of non-lytic predictions. The corresponding fractions were 28.5% and 12.3% for ZymCTRL and 73.8% and 46.1% for the ambiguous structural collection. The classifier and descriptor-space geometry therefore captured related but non-equivalent properties of the sequences.

The t-SNE projections illustrate the same relationships without defining them quantitatively (Figure **4**C–E). Prediction Extension sequences largely overlapped the Endolysin-dominated reference region. ZymCTRL sequences formed several peripheral structures with more limited overlap. The ambiguous structural collection showed clearer subtype-associated organization, including a prominent Depolymerase-like population. All quantitative support criteria remained defined in the complete descriptor space.

Subtype-specific analyses exposed several cases in which prediction and reference support diverged. The clearest example involved ZymCTRL VAL assignments. None of the 133 predicted VAL sequences fell within the VAL-specific reference coverage or satisfied the local geometry criteria. Their physicochemical profiles were also strongly displaced from the annotated VAL population. The classifier therefore detected features associated with its learned VAL decision region, while the descriptor-based reference analysis provided no comparable support from the available reference space.

The opposite pattern was observed for ambiguous structural Depolymerases (Figure **4**F). Property support was observed for 90.1% of these predictions, subtype-specific coverage for 91.2%, and local geometric support for 74.8%. Full concordance reached 65.9%. Prediction Extension Endolysins also showed substantial support. The corresponding fractions were 85.7% for physicochemical support, 60.0% for subtype coverage, and 92.4% for local geometry, resulting in 54.2% full concordance.

**Figure 4.**
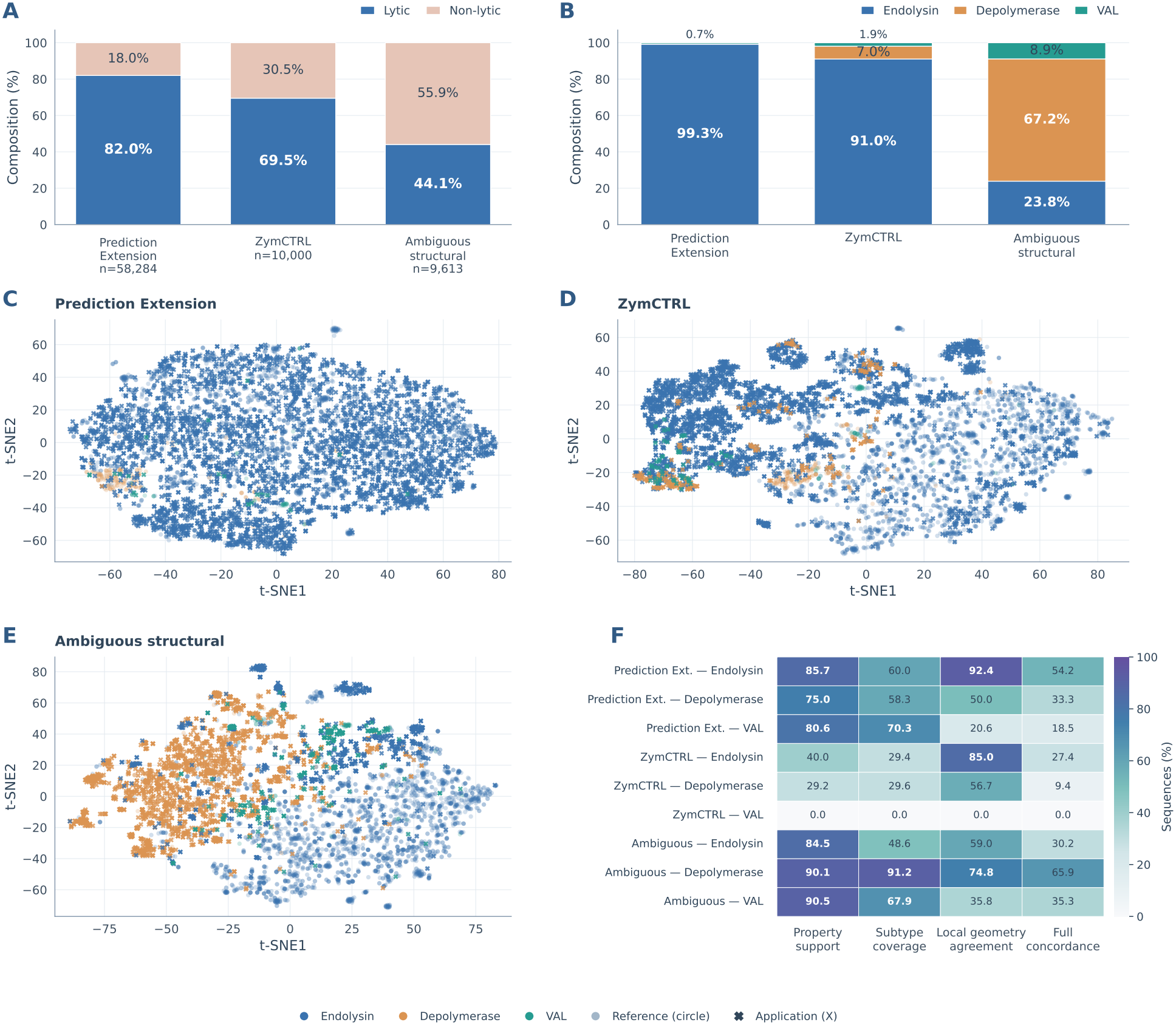
Application and reference-space contextualization of the frozen PhageLysClass architecture. **A**, fraction of sequences classified as globally lytic in the PhageLys Prediction Extension, ZymCTRL-generated collection, and ambiguous structural challenge set. **B**, functional composition of globally lytic predictions assigned to Endolysin, Depolymerase, or VAL. **C–E**, t-SNE visualization of reference proteins and application sequences for the Prediction Extension, ZymCTRL, and ambiguous structural collections. **F**, integrated support for each source and predicted subtype. Property support describes compatibility with subtype-specific physicochemical reference intervals. Subtype coverage describes membership within the corresponding reference-distance boundary. Local geometry agreement summarizes neighborhood-based sub-type support. Full concordance identifies sequences satisfying all three criteria. Quantitative support and distance calculations were performed in the complete standardized 26-dimensional descriptor space.

Support was lower for minority classes and generated sequences. Full concordance reached 33.3% for Prediction Extension Depolymerases and 18.5% for Prediction Extension VALs. Among ZymCTRL predictions, 27.4% of Endolysins and 9.4% of Depolymerases satisfied all three contextualization criteria. No predicted ZymCTRL VAL satisfied any of the three support axes.

These analyses distinguish prediction from reference support. The Prediction Extension primarily expanded an Endolysin-like region already represented by evidence-supported proteins. ZymCTRL showed that generated sequences can satisfy learned model boundaries while remaining distant from natural physicochemical reference space. The ambiguous structural challenge identified a Depolymerase-rich subset showing concordance between model predictions and complementary physicochemical and geometric reference-space criteria. Discordant sequences were retained and were not reassigned by the contextualization layer.

### 3.4 Scope, limitations, and recommended use of PhageLysClass

PhageLysClass was developed to recognize sequence patterns associated with phage lytic and surface-degrading proteins under controlled changes in biological task definition and sequence-space relatedness. Its predictions should therefore be interpreted within the evidence structure represented during model development. A positive global prediction indicates compatibility with the learned lytic decision space. It does not demonstrate enzymatic activity, substrate specificity, antibacterial activity, or biological function in a particular phage context.

Evidence support differs substantially among the three principal functional classes. Endolysins dominate the positive reference universe, while Depolymerases and especially VALs are represented by much smaller sequence populations. This imbalance was reflected in both the benchmark and the external applications. VAL showed the greatest variability under sequence-space restriction and the weakest agreement in several external contextualization analyses. High discrimination metrics obtained for VAL under individual test conditions therefore should not be interpreted as equivalent to the evidence available for Endolysin or Depolymerase recognition.

The global and multiclass components also answer different questions. Global recognition evaluates compatibility with the evidence-supported lytic population relative to the explicit non-target background. The multiclass component assigns a functional subtype conditional on a sequence first being classified as lytic. Auxiliary subtype models provide additional compatibility scores and do not replace either primary decision. These outputs should remain separated in downstream use.

The application collections do not constitute conventional external validation datasets. The Prediction Extension contains computationally annotated proteins derived from the broader PhageLysData universe. The ambiguous structural set was intentionally constructed from proteins with uncertain functional relationships to lytic and surface-degrading activities. ZymCTRL sequences were generated under enzyme-specific conditioning and do not provide experimentally established functional labels. Their value lies in probing how the frozen model behaves outside the sequence populations used during development.

Reference-space contextualization addresses a separate limitation. A high model score can occur for proteins whose physicochemical properties differ markedly from those of evidence-supported reference sequences. This behaviour was most evident among ZymCTRL proteins. Conversely, some predictions from the ambiguous structural set showed strong agreement with complementary physicochemical and geometric reference-space criteria. These criteria were evaluated independently of the ESM2-T30 classifier but were derived from the same 26 sequence descriptors. Their agreement therefore provides complementary characterization of reference compatibility rather than statistically independent evidence of biological function. Reference support should be interpreted as an additional evidence layer and not as confirmation of molecular function. The contextualization boundaries also depend on the current composition of PhageLysData and will change as experimentally supported sequence diversity increases.

PhageLysClass is best suited to candidate prioritization within a broader protein-analysis workflow. Sequences supported by global prediction, conditional subtype assignment, and compatible reference-space properties represent stronger candidates for downstream analysis than sequences supported only by the classifier. Discordant predictions remain informative because they identify extrapolative regions where the model detects a learned functional signal outside the physicochemical space currently represented by annotated proteins.

Downstream prioritization should incorporate evidence not used directly by the classifier. Domain architecture can determine whether a predicted catalytic activity is compatible with the organization of the complete protein. Structural analysis can distinguish catalytic modules from related non-catalytic folds and can clarify the role of virion-associated proteins. Genomic context can establish whether a candidate occurs in a biologically plausible phage lysis, adsorption, or structural module. Taxonomic information and available experimental annotations can further constrain interpretation. Experimental characterization remains necessary to establish enzymatic activity and biological function.

The framework therefore supports a staged use model in which sequence-based prediction identifies candidates, subtype classification defines a functional hypothesis, and physicochemical reference-space support, together with additional structural, genomic, and experimental evidence, guides how strongly that hypothesis should be pursued. This separation is especially important for generated proteins and poorly annotated phage proteins, where model confidence alone cannot establish biological plausibility.

## 4 Conclusions

PhageLysClass shows that sequence-based recognition of phage lytic and surface-degrading proteins depends as strongly on biological task definition and sequence-space control as on model choice. The evidence universe was dominated by unresolved proteins, while experimentally supported classes differed markedly in sequence architecture, physicochemical properties, and representation-space organization. These differences were also background dependent, emphasizing the importance of separating explicit non-target proteins from sequences with unresolved function.

Matched evaluation across redundancy and train–test homology regimes identified ESM2-T30 as the most transferable representation and showed that redundancy control exerted a stronger effect on generalization than homology separation alone. The final architecture therefore emerged from robustness across sequence-space conditions instead of from a single optimal benchmark result. Under the strict R30/S30 condition, the global lytic-recognition component retained an MCC of approximately 0.89, while the multiclass component remained highly accurate for Endolysin and Depolymerase assignment.

Application of the frozen system further showed that model prediction and biological reference support are distinct quantities. Prediction Extension proteins largely remained within evidence-supported sequence-property space, generated ZymCTRL proteins frequently represented extrapolative predictions, and ambiguous structural proteins contained a strongly supported Depolymerase-like subset, with agreement between model assignments and complementary physicochemical and geometric reference-space criteria. PhageLysClass should therefore be used as an evidence-aware prioritization framework in which predictive assignments are interpreted together with physicochemical, structural, genomic, and experimental information. This design provides a controlled basis for expanding phage lytic-protein discovery while preserving uncertainty in poorly characterized regions of protein sequence space.

## Author contributions statement

AO-N, JR, and DM-O: conceptualization. DM-O: methodology. DM-O: validation. AO-N, JR, MEL, AV and DM-O: investigation. AO-N, JR, MEL and DM-O: writing and editing. JR and MEL: supervision and funding resources. JR: project administration. All authors reviewed and approved the final version of the manuscript.

## Data and Code Availability

The research datasets, numerical protein representations, redundancy-controlled partitions, benchmark results, individual prediction outputs, and pretrained models supporting this study are publicly available through the PhageLysClass v1.0 Zenodo archive (https://doi.org/10.5281/zenodo.24354677). The source code, computational workflows, analysis notebooks, and inference tools are available in the PhageLysClass GitHub repository (https://github.com/kren-ai-lab/phagelysclass). The archived resources preserve frozen task definitions, experimental configurations, sequence identifiers, and computational outputs to support reproducibility and independent reuse. Software versions, model configurations, computational environments, random seeds, and provenance controls are documented in Supplementary Methods Section S1.8.

## Acknowledgments

This research was funded by the Chilean National Agency for Research and Development (ANID), through FONDECYT Exploración Grant No. 13240071.

## AI Statement

As non-native English speakers, the authors used ChatGPT (OpenAI, GPT-5.6 Thinking; accessed July 2026) for English editing, clarity, and consistency. The authors critically reviewed and edited all outputs, verified the final text, and take full responsibility for the manuscript.

