## Supplementary Material for "PhageLysClass: Evidence-aware classification of phage lytic proteins using protein language models"

#### Supplementary contents

**S1 Supplementary Methods** **2**

### S1 Supplementary Methods

#### S1.1 Construction of the evidence-state universe

The PhageLysClass evidence universe was constructed from the frozen PhageLysData Core and a complementary UniProtKB phage-protein collection. The PhageLysData Core contained 11,867 unique exact-sequence entities and was the unique source of positive biological labels (Medina Ortiz et al., 2026). UniProtKB release 2026_02 (Bateman et al., 2025) was used to construct the complementary background through taxonomy-based retrieval, without using functional terms to predefine negative or unlabeled proteins.

UniProtKB retrieval comprised proteins assigned to Caudoviricetes (`taxonomy_id:2731619`) together with named phage or bacteriophage proteins from viral taxa outside Caudoviricetes. Reviewed and unreviewed records were retained. Retrieval used the UniProtKB REST search endpoint with cursor pagination and 500 records per page. The retained fields included accession and entry identifiers, protein and gene names, organism and taxonomic information, virus host, reviewed status, protein-existence evidence, annotation score, fragment status, amino-acid sequence, functional and catalytic-activity comments, EC numbers, GO identifiers, keywords, and record dates.

Sequences were normalized to uppercase after removal of whitespace, and exact amino-acid sequence was used as the biological identity unit. A SHA-256 sequence hash was retained as a stable identifier. UniProtKB accessions encoding the same sequence were collapsed while preserving their accession-level annotations and provenance. The principal sequence counts produced during universe construction are summarized in Table **S1**.

**Supplementary Table S1**: **Construction of the frozen PhageLysClass evidence universe.**

| Source or integration step | Records | Exact sequences | Role |
|---|---|---|---|
| PhageLysData Core | – | 11,867 | Evidence-supported positive reference and sole source of positive labels |
| UniProtKB phage retrieval | 570,195 | 437,447 | Taxonomy-derived background for evidence-state assignment |
| Exact Core–UniProtKB overlap | – | 6,985 | Decontamination overlap; represented only by the corresponding Core entity |
| Final integrated universe | – | 442,329 | Unique exact-sequence entities used for evidence-state construction |

UniProtKB sequences were classified only after retrieval using a frozen evidence hierarchy. Rules inspected protein names, catalytic annotations, EC assignments, keywords, and selected curated functional comments. Exact Core overlap had highest precedence. Remaining records were evaluated for direct lytic-related evidence, structural or adsorption-associated ambiguity, lysis-adjacent functions, explicit alternative functions, explicit unresolved annotation, and other unresolved status. Direct lytic identity was based on protein-identity and catalytic-evidence fields and was not inferred from unrestricted functional free text. Generic protease or peptidase terminology was also insufficient by itself to define an alternative non-lytic function.

The complete state hierarchy, precedence order, exact-sequence counts, and downstream roles are shown in Table **S2**. If different accessions encoding the same sequence supported different states, the highest-priority state was retained. Any target-like or structurally ambiguous annotation prevented the corresponding sequence from entering a negative or unlabeled population.

The Core contained 9,888 Endolysins, 712 Depolymerases, 241 virion-associated lytic enzymes (VALs), 999 proteins with multiple non-exclusive lytic assignments, and 27 proteins assigned to other phage-envelope lytic functions. All 11,867 Core proteins were valid positives for global lytic-recognition tasks. Only the 10,841 single-class Endolysin, Depolymerase, and VAL proteins were eligible for the clean multiclass formulation.

**Supplementary Table S2**: **Frozen evidence-state hierarchy of the PhageLysClass universe.**

| Priority | State | Sequences | Operational evidence | Downstream role |
|---|---|---|---|---|
| 1 | P_core | 11,867 | Evidence-supported exact-sequence entity from the PhageLysData Core | Positive reference |
| 2 | challenge_lytic_related | 319 | Direct lytic-like UniProtKB evidence without exact Core overlap | Target-like challenge population |
| 3 | challenge_ambiguous_struct. | 10,885 | Tail-, adsorption-, receptor-binding-, baseplate-, or related structural evidence potentially overlapping VAL or Depolymerase function | Structural challenge population |
| 4 | N_hard | 4,327 | Lysis-adjacent proteins, including holins, antiholins, spanins, and related generally non-enzymatic functions | Hard supervised background |
| 5 | N_easy | 54,711 | Explicit alternative phage functions without higher-priority target-like evidence | Alternative-function supervised background |
| 6 | U_explicit | 228,075 | Explicitly hypothetical, uncharacterized, DUF/UPF-associated, or low-information annotation | Conservative unlabeled population |
| 7 | U_other_unresolved | 132,145 | Remaining proteins that could not be assigned safely to another evidence state | Additional broad unlabeled population |
| **Total** | | **442,329** | | |

**Supplementary Table S3**: **Frozen PhageLysClass prediction-task definitions before redundancy and homology control.**

| Task | Positive or target population | Comparison population | Total | Purpose |
|---|---|---|---|---|
| Global easy | Core, $n = 11{,}867$ | N_easy, $n = 54{,}711$ | 66,578 | Alternative-function discrimination |
| Global hard | Core, $n = 11{,}867$ | N_hard, $n = 4{,}327$ | 16,194 | Lysis-adjacent discrimination |
| Global mixed | Core, $n = 11{,}867$ | N_easy + N_hard, $n = 59{,}038$ | 70,905 | Primary global lytic-recognition task |
| Endolysin binary | Endolysin, $n = 9{,}888$ | N_easy + N_hard, $n = 59{,}038$ | 68,926 | Subtype compatibility |
| Depolymerase binary | Depolymerase, $n = 712$ | N_easy + N_hard, $n = 59{,}038$ | 59,750 | Subtype compatibility |
| VAL binary | VAL, $n = 241$ | N_easy + N_hard, $n = 59{,}038$ | 59,279 | Subtype compatibility |
| Multiclass clean | Endolysin / Depolymerase / VAL | 9,888 / 712 / 241 | 10,841 | Primary functional assignment |
| PU conservative | Core, $n = 11{,}867$ | U_explicit, $n = 228{,}075$ | 239,942 | PU learning with explicit unresolved annotation |
| PU broad | Core, $n = 11{,}867$ | U_explicit + U_other_unresolved, $n = 360{,}220$ | 372,087 | PU learning across the broader unresolved universe |

The evidence hierarchy was frozen after an annotation audit covering state composition, conflicting rule assignments, annotation-quality metadata, Core overlaps, and reproducible random samples of 75 proteins from each evidence state using random seed 42. The audit identified and corrected cases in which unrestricted functional text could propagate lytic terminology to non-enzymatic lysis proteins and cases in which broad protease terminology could produce inappropriate alternative-function assignments. No evidence state was changed after inspection of predictive performance.

Prediction tasks were then materialized directly from the frozen evidence states. No redundancy reduction, homology filtering, class balancing, representation generation, or model fitting was performed during task definition. The nine resulting supervised and positive–unlabeled formulations are summarized in Table **S3**.

The three global supervised tasks shared the complete positive Core and differed only in their comparison population. The three subtype binary tasks used the same combined explicit non-lytic background but excluded all other lytic classes from the negative population. The clean multiclass task contained only single-class Endolysin, Depolymerase, and VAL proteins. The two PU tasks contained no explicit negative proteins; their unlabeled sequences retained unknown biological status throughout model development.

The UniProtKB lytic-related and ambiguous structural states were excluded from all supervised and PU fitting. They were preserved as challenge populations for subsequent application analyses. Core proteins with multiple or other lytic assignments remained positive examples in global recognition but were excluded from subtype-specific and multiclass fitting because they did not provide an unambiguous single-class label.

All evidence states and task memberships were frozen at this point. Subsequent redundancy reduction, homology-aware partitioning, predictive-eligibility filtering, representation generation, and model optimization operated on these predefined datasets and were not permitted to redefine biological state or task membership.

##### S1.2 Protein characterization and representation-space analysis

Protein characterization was performed on the frozen universe of 442,329 unique sequences before redundancy reduction or model development. Sequence properties were calculated using Roxy v0.2.0 (Medina-Ortiz et al., 2026a), and protein language model embeddings were generated with Sylphy (Medina-Ortiz et al., 2026b). The five characterization spaces are summarized in Table **S4**.

**Supplementary Table S4**: **Sequence-property and protein language model spaces used for characterization.**

| Space | Source | Dimension | Distance | Definition |
|---|---|---|---|---|
| Sequence composition | Roxy | 71 | Euclidean | Amino-acid frequencies, sequence complexity, compositional bias, and grouped residue properties |
| Physicochemical | Roxy | 27 | Euclidean | Global physicochemical, charge, order/disorder, structural-propensity, and AAindex descriptors |
| Combined explicit | Roxy | 98 | Euclidean | Concatenation of composition and physicochemical descriptors |
| ESM2-T6 8M | Sylphy | 320 | Cosine | Mean-pooled final hidden-layer embeddings |
| ESM2-T30 150M | Sylphy | 640 | Cosine | Mean-pooled final hidden-layer embeddings |

The initial composition panel contained 122 variables derived from the Roxy `aac`, `entropy_complexity`, `compositional_bias`, and `grouped` modules. Removal of raw counts, redundant size-dependent variables, incomplete descriptors, and non-variable features yielded 71 variables for multivariate analysis. The physicochemical panel contained 27 descriptors from the `global_basic`, `charge`, `order_disorder`, `structural_propensity`, and `aaindex` modules (Kawashima et al., 2007). Molecular weight was retained for initial characterization but excluded from the subsequent post-prediction panel, leaving 26 descriptors. These characterization spaces were distinct from the 48-feature Roxy representation used for predictive benchmarking.

ESM2 analysis required canonical amino-acid sequences of at most 1,024 residues. Both models represented the same 434,768 eligible proteins. The statistical comparisons, multivariate analyses, and visualization parameters are summarized in Table **S5**.

Non-finite descriptors were excluded from the corresponding multivariate feature spaces without imputing values or removing protein sequences. Sequence length was analyzed separately from the physicochemical effect-size rankings. Cross-representation comparisons used matched protein populations, with Euclidean distances for standardized explicit descriptors and cosine distances for ESM2 embeddings. All analyses were descriptive and did not influence evidence-state assignment, dataset partitioning, or predictive model selection.

**Supplementary Table S5: Statistical and geometric analysis configurations.**

| Analysis | Configuration |
|---|---|
| Biological comparisons | `P_core` versus `N_easy`, `N_hard`, and `N_mixed`; pairwise comparisons among clean Endolysin, Depolymerase, and VAL references |
| Univariate statistics | Mann–Whitney *U* test, Cliff's delta, and Benjamini–Hochberg correction; Kruskal–Wallis omnibus tests for physicochemical descriptors |
| Explicit-space geometry | StandardScaler; Euclidean distance; silhouette score, within-group dispersion, centroid distances, and same-label nearest-neighbor fraction |
| ESM2-space geometry | Cosine distance; silhouette score and same-label nearest-neighbor fraction |
| Neighborhood analysis | $k = 10$, 25, and 50; $k = 25$ for matched cross-representation comparisons |
| Balanced repetitions | Explicit spaces: 10 global and 20 subtype repeats; ESM2 spaces: 5 global and 10 subtype repeats |
| Matched representation comparison | Common set of 434,768 PLM-eligible proteins; five balanced repeats of 1,500 proteins per class for each Core–background contrast; silhouette and macro-averaged same-label neighbor fraction at $k = 25$ |
| Explicit-space projections | PCA: up to 10 components; UMAP: Euclidean distance, 30 neighbors, minimum distance 0.2, seed 42 |
| ESM2 projections | PCA: 20 components; UMAP: cosine distance, 30 neighbors, minimum distance 0.2, seed 42; t-SNE: PCA to 50 components, perplexity 30, 1,000 iterations, seed 42 |

#### S1.3 Redundancy control and homology-aware partitioning

Sequence redundancy and train–test homology were controlled sequentially using BioSieve and MMseqs2 (Villegas et al., 2026; Steinegger and Söding, 2017). Four redundancy regimes (R90, R70, R50, and R30) and four homology-aware splitting regimes (S90, S70, S50, and S30) were evaluated. The complete configuration is summarized in Table **S6**.

**Supplementary Table S6: Sequence redundancy and homology-aware partitioning used in PhageLysClass.**

| Component | Regimes | Scope | Configuration |
|---|---|---|---|
| Redundancy control | R90, R70, R50, R30 | Applied independently to each task and separately within its biological groups | MMseqs2-based reduction at 90%, 70%, 50%, or 30% sequence identity; each R dataset generated directly from the original frozen task |
| Homology clustering | S90, S70, S50, S30 | Applied to the complete redundancy-controlled task after recombination of biological groups | MMseqs2 minimum sequence identity of 90%, 70%, 50%, or 30%; minimum sequence coverage 80%; coverage mode 0; cluster mode 0 |
| Outer partition | Train / test | Whole MMseqs2 clusters | Approximately 70% training and 30% held-out test; no validation partition generated at this stage |
| Cluster integrity | All S conditions | Train–test boundary | All members of an MMseqs2 cluster required to remain in the same partition |
| Factorial design | 4 R × 4 S | Each prediction task | 16 outer conditions per task; 144 conditions across the nine supervised and positive–unlabeled tasks |

Redundancy reduction was performed separately within the biological groups defining each task before the retained representatives were recombined. Global binary, subtype-specific, and positive–unlabeled tasks were reduced within their corresponding positive, negative, or unlabeled populations. The clean multiclass task was reduced independently within the Endolysin, Depolymerase, and VAL classes. This prevented a representative from one biological group from replacing a sequence belonging to another group. Homology between different groups was not removed during redundancy reduction.

Each R dataset was subsequently partitioned independently under S90, S70, S50, and S30. Unlike redundancy reduction, MMseqs2 (Steinegger and Söding, 2017) clustering for S was performed across the complete task so that homologous proteins from different biological groups could not be separated between train and test. Whole clusters were assigned to one partition, with approximately 30% of sequences targeted for the held-out test set. Cluster integrity had priority over obtaining an exact test fraction.

A small number of initial BioSieve assignments contained members of the same precomputed cluster in both partitions. These cases were corrected by assigning the complete cluster to the partition that produced a test size closest to the 30% target. If both alternatives produced the same deviation, the cluster was assigned to the partition containing the majority of its members in the initial split. MMseqs2 cluster membership itself was not modified.

Every final R–S condition was audited before model development. The audit required complete and unique assignment of all input sequences, absence of exact-sequence overlap between train and test, and absence of MMseqs2 clusters spanning both partitions. Sequence counts, retained fractions, train/test composition, and cluster counts were recorded for each condition.

Challenge populations were not subjected to the R–S procedure. Protein representations, model predictions, and downstream performance were not used during redundancy reduction or partition construction.

##### S1.4 Model optimization and positive–unlabeled learning

###### S1.4.1 Representation eligibility and preprocessing

A common predictive universe was defined before model fitting so that representation comparisons used the same proteins within each task and $R \times S$ condition. Eligible proteins were required to belong to the frozen exact-sequence universe, contain only the 20 canonical amino acids, and have a sequence length of at most 1,024 residues. This filtering preserved the outer train–test assignments and the homology-cluster identifiers defined in S1.3. Representation availability was evaluated only after this biological eligibility filter. If an eligible sequence was missing from a representation, the corresponding representation was considered incomplete rather than removing the protein from the benchmark.

The predictive benchmark evaluated four protein representations, including 48 sequence-derived descriptors calculated with Roxy, fixed-length one-hot encoding, and ESM2-T6 and ESM2-T30 embeddings. Their dimensions and preprocessing configurations are summarized in Table **S7**.

**Supplementary Table S7: Protein representations used in the predictive benchmark.**

| Representation | Type | Dimension | Predictive preprocessing |
|---|---|---|---|
| Roxy | Sequence descriptors | 48 | Standardized for logistic regression, $k$-nearest neighbors, and multilayer perceptron models; used without scaling for random forest and XGBoost |
| One-hot | Sequence encoding | 20,480 | Sparse encoding of 1,024 positions and 20 canonical amino acids; used without additional scaling or dimensionality reduction |
| ESM2-T6 | Protein language model | 320 | Final-layer mean-pooled FP32 embedding generated with Sylphy; standardized for logistic regression, $k$-nearest neighbors, and multilayer perceptron models |
| ESM2-T30 | Protein language model | 640 | Final-layer mean-pooled FP32 embedding generated with Sylphy; standardized for logistic regression, $k$-nearest neighbors, and multilayer perceptron models |

###### S1.4.2 Learner configuration, hyperparameter spaces, and group-aware inner validation

Five supervised learners were evaluated for every representation and $R \times S$ condition, including logistic regression (LR), random forest (RF), $k$-nearest neighbors (KNN), XGBoost (XGB), and a multilayer perceptron (MLP). Instead of unrestricted hyperparameter optimization, three predefined configurations of increasing or alternative model complexity were evaluated for each learner. The complete configuration panel is shown in Table **S8**.

Class imbalance was handled within the fitting procedure. LR used balanced class weights, RF used balanced subsample weights, and XGBoost received sample weights calculated from the class frequencies of the current fitting partition. No additional over- or undersampling was applied to the supervised tasks.

Model configuration was selected using a fixed inner validation partition generated from the outer training set of each task and $R \times S$ condition. Five candidate partitions were generated with stratified group five-fold splitting using the S-level homology clusters as groups and random seed 42. Complete homology clusters

**Supplementary Table S8: Predefined supervised-learning configurations.**

| Learner | Configuration | Variable parameters | Fixed settings |
|---|---|---|---|
| LR | LR01 / LR02 / LR03 | $C = 0.1$ / 1.0 / 10.0 | L2 regularization, `lbfgs`, maximum 2,000 iterations, balanced class weights |
| RF | RF01 | 300 trees; unrestricted depth; minimum leaf size 1 | Square-root feature sampling; balanced subsample weights |
| RF | RF02 | 500 trees; unrestricted depth; minimum leaf size 2 | Square-root feature sampling; balanced subsample weights |
| RF | RF03 | 500 trees; maximum depth 30; minimum leaf size 1 | Square-root feature sampling; balanced subsample weights |
| KNN | KNN01 / KNN02 / KNN03 | $k = 3$ / 5 / 11 | Cosine distance, distance-weighted voting, brute-force neighbor search |
| XGB | XGB01 | 300 trees; depth 4; learning rate 0.10 | Subsample 0.8; column subsample 0.8; histogram tree method |
| XGB | XGB02 | 500 trees; depth 6; learning rate 0.05 | Subsample 0.8; column subsample 0.8; histogram tree method |
| XGB | XGB03 | 800 trees; depth 4; learning rate 0.03 | Subsample 0.8; column subsample 0.8; histogram tree method |
| MLP | MLP01 | 256 hidden units; $\alpha = 10^{-4}$; learning rate $10^{-3}$ | ReLU activation, Adam optimizer, batch size 256, maximum 300 iterations |
| MLP | MLP02 | 512 and 128 hidden units; $\alpha = 10^{-4}$; learning rate $5 \times 10^{-4}$ | ReLU activation, Adam optimizer, batch size 256, maximum 300 iterations |
| MLP | MLP03 | 256, 128, and 64 hidden units; $\alpha = 10^{-3}$; learning rate $5 \times 10^{-4}$ | ReLU activation, Adam optimizer, batch size 256, maximum 300 iterations |

were therefore kept within either inner training or validation. Among the five candidate validation folds, the partition closest to the target validation fraction of 20% while preserving the overall class composition was selected. Candidate partitions that did not retain all observed classes in both inner training and validation were rejected. The resulting inner assignment was frozen and reused across all representations, learners, and configurations within the same task and $R \times S$ condition.

Each of the three predefined configurations was fitted on the inner-training partition and evaluated on the same validation proteins. Binary configurations were ranked first by Matthews correlation coefficient (MCC) and then by average precision (AP). Multiclass configurations were ranked first by multiclass MCC and then by macro-averaged F1 score. Configuration order was used only to resolve remaining ties. No information from the outer test partition was used during configuration selection.

After selection, the chosen configuration was rebuilt using the complete eligible outer-training partition and evaluated once on the frozen outer test set. Binary probability-based decisions used a threshold of 0.5; no task-specific threshold optimization was performed. The final supervised benchmark therefore comprised seven supervised tasks, 16 $R \times S$ conditions, four representations, and five learners, corresponding to 2,240 matched experiment families.

##### S1.4.3 Positive–unlabeled learning configurations and evaluation

Positive–unlabeled learning was evaluated separately for the conservative and broad PU task definitions described in S1.1. The final PU benchmark used the same four predictive representations and all 16 $R \times S$ conditions, but deliberately restricted the base learners to LR and XGBoost. Fixed intermediate configurations, LR02 and XGB02, were used throughout the PU analysis. No hyperparameter selection was performed using unlabeled proteins.

Three PU strategies were evaluated, as summarized in Table **S9**. In all cases, the observed label distinguished known positives (P) from unlabeled proteins (U); U was not interpreted as a confirmed biological negative class.

The naive P-versus-U model and BaggingPU were fitted directly on the complete eligible outer-training partition because their learner configurations were fixed in advance. For BaggingPU, the unlabeled subset was sampled independently for each of the 30 bags using random seed 42 and a U:P sampling ratio of 1:1. The mean probability across bags was used as the final score.

**Supplementary Table S9: Positive–unlabeled learning configurations used in the final benchmark.**

| PU strategy | Base learners | Configuration |
|---|---|---|
| Naive P-versus-U | LR02, XGB02 | Known positives were assigned the observed label 1 and unlabeled proteins the observed label 0. Models were fitted on the complete eligible outer-training partition and used as a diagnostic P-versus-U ranking baseline. |
| BaggingPU | LR02, XGB02 | Thirty bags were fitted. Each bag contained all available training positives and an equally sized subset of unlabeled proteins sampled without replacement. Test scores were calculated as the mean positive probability across the 30 models, and the per-protein standard deviation was retained. |
| Elkan–Noto | LR02, XGB02 | The labeling propensity $c$ was estimated by five-fold stratified group cross-fitting within the outer-training partition, using S-level homology clusters as groups. The final base learner was then fitted on the complete outer-training partition and corrected scores were calculated as the raw positive score divided by $c$, clipped to the interval $[0, 1]$ (Nanduri et al., 2026). |

For Elkan–Noto learning, the labeling propensity $c$ was estimated without scoring positives used to fit the corresponding model (Jain et al., 2016). Five stratified group folds were generated within the outer-training partition using the S-level homology clusters and random seed 42. For each fold, positive scores were obtained only for held-out known positives, and $c$ was calculated as the mean of these out-of-fold positive scores. A final model was then fitted on the complete outer-training partition. Both the raw score and the corrected score were retained, together with the fraction of corrected probabilities that required clipping above 1.

PU evaluation focused on separation and ranking of known positives relative to the unlabeled population. For each held-out outer test set, the analysis recorded the median scores of P and U proteins, the 90th, 95th, and 99th percentiles of the U-score distribution, and the fraction of known positives scoring above each of these U thresholds. The fractions of P and U proteins with scores of at least 0.5 were also recorded. These quantities describe recovery of known positives and ranking within the unlabeled population; they were not interpreted as biological sensitivity, specificity, or negative-class accuracy.

The final PU design comprised two PU universes, 16 $R \times S$ conditions, four representations, two base learners, and three PU strategies, producing 768 PU experiment families. The PU analysis remained separate from the supervised benchmark and did not alter task definitions, representation selection, or supervised model configuration.

##### S1.5 Generalization analysis and predictive architecture selection

Generalization analysis was performed on the frozen outer-test predictions generated for the complete supervised benchmark. For binary tasks, MCC was used as the primary performance measure,

$$\mathrm{MCC} = \frac{TP\,TN - FP\,FN}{\sqrt{(TP + FP)(TP + FN)(TN + FP)(TN + FN)}},$$

with average precision (AP), ROC-AUC, balanced accuracy, F1 score, sensitivity, and specificity used as complementary measures. Because AP depends directly on class prevalence, comparisons between task formulations with different positive fractions also used prevalence-normalized AP skill,

$$\mathrm{AP}_{\mathrm{skill}} = \frac{\mathrm{AP} - \pi}{1 - \pi},$$

where $\pi$ is the positive prevalence in the evaluated test set. A value of zero therefore corresponds to the prevalence baseline and a value of one to perfect ranking. Multiclass performance was summarized primarily using multiclass MCC and macro-averaged F1 score, with class-specific precision, recall, F1 score, and support retained for interpretation.

Comparisons between representations or learners within the same task and $R \times S$ condition were matched because all models were evaluated on the same frozen test proteins. For two strategies $A$ and $B$, the performance contrast was defined as

$$\Delta M = M_A - M_B,$$

where $M$ denotes the metric under consideration. Comparisons across redundancy or homology conditions were not assumed to be paired because the corresponding outer-test populations could differ. These native-test contrasts were therefore complemented by common-test analyses. For two conditions $a$ and $b$, the shared evaluation population was defined as

$$\mathcal{T}_{a,b} = \mathcal{T}_a \cap \mathcal{T}_b,$$

and both models were re-evaluated using only the proteins in $\mathcal{T}_{a,b}$. The corresponding common-test contrast was

$$\Delta M_{\text{common}} = M_A(\mathcal{T}_{a,b}) - M_B(\mathcal{T}_{a,b}).$$

This analysis was used to distinguish changes in prediction behavior from changes caused by different test-set composition.

Uncertainty was quantified for the principal matched comparisons using 1,000 paired bootstrap replicates. Proteins were resampled with replacement while preserving the observed class structure, and the same resampled entities were used for both members of each comparison. Multiclass resampling was performed within class. For bootstrap replicate $b$,

$$\Delta M^{(b)} = M_A^{(b)} - M_B^{(b)}.$$

The 95% confidence interval was obtained from the 2.5th and 97.5th percentiles of the bootstrap distribution. Directional consistency was defined as

$$C_{\text{dir}} = \max \left[ P\left(\Delta M^{(b)} > 0\right), P\left(\Delta M^{(b)} < 0\right)\right].$$

Bootstrap analysis was restricted to scientifically relevant paired contrasts, including selected representation, learner, and common-test $R/S$ comparisons; it was not applied exhaustively to every possible pair in the factorial benchmark.

Model robustness was evaluated across the complete set of 16 $R \times S$ conditions for each task–representation–learner strategy. If $M_{j,r,s}$ denotes the MCC of strategy $j$ under redundancy regime $r$ and homology regime $s$, its average, worst-case, strict-condition, and range statistics were defined as

$$\overline{M}_j = \frac{1}{16} \sum_{r,s} M_{j,r,s},$$

$$M_j^{\text{worst}} = \min_{r,s} M_{j,r,s},$$

$$M_j^{\text{strict}} = M_{j,30,30},$$

and

$$G_j = \max_{r,s} M_{j,r,s} - \min_{r,s} M_{j,r,s}.$$

Rank-based robustness was evaluated in parallel. For each $R \times S$ cell, strategies were ranked within the corresponding task and the regret of strategy $j$ was defined as

$$g_{j,r,s} = \max_k M_{k,r,s} - M_{j,r,s}.$$

Mean rank and mean regret were then calculated across the 16 cells. Additional summaries included performance under $S30$, performance under $R30$, sensitivity to changes in R and S, and the magnitude of the $R \times S$ interaction. These quantities were kept separate instead of being combined into a single robustness score.

Individual models advanced to ensemble analysis through a multi-criterion portfolio procedure. A strategy was eligible if it either led an explicit operational criterion or had a mean MCC within 0.02 of the best mean MCC for that task,

$$\overline{M}_j \geq \overline{M}_{\max} - 0.02.$$

Operational criteria included average performance, worst-case performance, strict $R30/S30$ performance, computational efficiency among competitive models, rank consistency, regret, remote-homology performance, strict-redundancy performance, and interaction sensitivity. Between two and four individual strategies were retained per prediction endpoint. No weighted sum of performance, robustness, uncertainty, and computational cost was used for candidate selection.

All non-singleton combinations of the retained individual strategies were subsequently evaluated as probability-level ensembles. For an ensemble containing $K$ binary classifiers, unweighted soft voting was defined as

$$p_{\text{ens}}(x) = \frac{1}{K} \sum_{k=1}^{K} p_k(x),$$

with a final positive decision at $p_{\text{ens}} \geq 0.5$. For multiclass prediction,

$$p_{\text{ens}}(c \mid x) = \frac{1}{K} \sum_{k=1}^{K} p_k(c \mid x),$$

and the predicted class was the class with maximum ensemble probability. Candidate predictions were aligned by exact sequence identity before aggregation, and ensemble performance was recalculated independently for all 16 $R \times S$ conditions.

Validation-weighted soft voting was evaluated as a secondary formulation. Member weights were based only on the selected inner-validation MCC,

$$w_k = \frac{\max(0, M_k^{\text{val}})}{\sum_j \max(0, M_j^{\text{val}})},$$

and ensemble probabilities were calculated as

$$p_{\text{wens}}(x) = \sum_{k=1}^{K} w_k p_k(x).$$

Outer-test performance was never used to determine ensemble weights or decision thresholds. Because validation-weighted voting did not provide a material advantage over uniform averaging, unweighted soft voting was frozen for production ensembles.

Final recipe selection remained endpoint-specific. Candidate recipes first had to fall within the competitive mean-performance band. Global lytic recognition then prioritized worst-case MCC, followed by strict $R30/S30$ and mean performance. Clean multiclass prediction prioritized strict-condition MCC, followed by worst-case and mean performance. Endolysin compatibility prioritized mean, worst-case, and strict performance, whereas Depolymerase compatibility prioritized strict, mean, and worst-case performance. Lower surface range and lower model complexity were used as additional tie-breaking criteria where appropriate. VAL was retained as a single-model auxiliary output because ensemble averaging introduced an unfavorable trade-off between average and strict-condition performance.

Three system organizations were subsequently considered: two core modules alone, five co-equal predictive modules, and two core modules with three auxiliary compatibility modules. Architecture selection was performed by constraint satisfaction rather than by an additional numerical score. The required constraints were an unambiguous primary inference path, broad global lytic recognition, primary multiclass functional assignment, preservation of subtype compatibility information, explicit treatment of VAL as evidence limited, separation of binary compatibility from multiclass functional assignment, a minimal default inference path, and compatibility with the frozen unweighted-ensemble policy. The resulting production organization comprised two core modules and three optional auxiliary modules. The frozen components are summarized in Table **S10**.

**Supplementary Table S10**: **Frozen PhageLysClass predictive architecture.**

| **Component** | **Task** | **Frozen learner composition** | **Role** |
|---|---|---|---|
| Global lytic core | Global mixed | KNN + MLP, unweighted soft voting | Primary lytic versus non-lytic/background decision |
| Functional core | Clean multiclass | XGBoost + LR, unweighted soft voting | Primary Endolysin, Depolymerase, or VAL assignment |
| Endolysin auxiliary | Endolysin binary | KNN + XGBoost + MLP, unweighted soft voting | Optional Endolysin compatibility score |
| Depolymerase auxiliary | Depolymerase binary | MLP + KNN, unweighted soft voting | Optional Depolymerase compatibility score |
| VAL auxiliary | VAL binary | MLP | Optional VAL compatibility score with evidence-limited interpretation |

The final architecture converged on ESM2-T30 for every retained component, allowing a single 640-dimensional embedding to be reused across the complete prediction stack. The default inference path therefore contains only the global lytic-recognition core followed, for proteins classified as lytic, by the clean multiclass functional core. The three subtype-specific binary components remain optional compatibility outputs and do not vote against or replace the multiclass prediction. The positive–unlabeled models were not incorporated into this architecture and remained a separate discovery-oriented workflow.

Architecture freezing fixed the task definitions, representation, learner membership, ensemble membership, voting procedure, component roles, and output precedence. The experimental R and S conditions were not converted into production hyperparameters. Instead, the configuration selected most frequently for each retained learner across the 16 validation-based $R \times S$ experiments was used as the production configuration, with validation performance used to resolve ties. Each retained estimator was then refitted using all eligible proteins from its corresponding frozen task definition, without preserving an outer benchmark split. This produced four fitted estimators in the default two-core path and ten fitted estimators when all auxiliary components were included. No application or challenge sequence was used during architecture selection or final refitting.

##### S1.6 Preparation and frozen inference of heterogeneous application collections

The frozen PhageLysClass architecture was applied to three collections excluded from model development, comprising the PhageLys Prediction Extension, ZymCTRL-generated proteins (Munsamy et al., 2022), and the UniProtKB ambiguous structural challenge population. Source-specific preparation yielded 77,897 proteins, as summarized in Table **S11**.

The PhageLys Prediction Extension comprised 745,092 unique sequences supported exclusively by prediction-derived annotations (Medina Ortiz et al., 2026). Proteins containing only the 20 canonical amino acids and no more than 1,024 residues were retained and subjected to MMseqs2 redundancy reduction at 30% sequence identity. This produced 58,284 representative sequences. Original identifiers, predicted functional annotations, and provenance information were preserved as metadata but were not used as classifier inputs.

ZymCTRL generation used the `AI4PD/ZymCTRL` checkpoint with `top_k`=9, repetition penalty 1.2, temperature 1.0, random seed 42, batch size 10, and maximum sequence length 1,024. Five conditioning EC classes represented peptidoglycan degradation (3.2.1.17, 3.2.1.52, 3.5.1.28, 4.2.2.29, and 3.4.24.75), and five represented surface-polysaccharide degradation (3.2.1.87, 3.2.1.129, 4.2.2.1, 4.2.2.3, and 3.2.1.65). Sequences were accepted only when generation terminated before the length limit, all residues were canonical, and the sequence was unique across the generated collection. Generation continued until 1,000 accepted sequences

**Supplementary Table S11**: **Application collections used with the frozen PhageLysClass architecture.**

| Collection | Origin | Principal preparation | Final sequences |
|---|---|---|---|
| PhageLys Prediction Extension | Prediction-only sequences from PhageLysData | Sequence-eligibility filtering and redundancy reduction at 30% identity | 58,284 |
| ZymCTRL | De novo proteins conditioned on ten enzyme classes | Canonical residues, generation completion, length, and uniqueness filters | 10,000 |
| Ambiguous structural challenge | UniProtKB proteins with tail-, adsorption-, receptor-binding-, or related structural annotations | Sequence-eligibility filtering and exact-sequence overlap audit | 9,613 |

were obtained per EC class. Perplexity was recorded but was not used for filtering. Conditioning EC classes were retained as metadata and were not treated as ground-truth functional subtype labels.

The ambiguous structural challenge comprised 10,885 unique UniProtKB proteins with tailspike, tail-fiber, receptor-binding, baseplate, adsorption-associated, or related annotations. These proteins were excluded from supervised and positive–unlabeled model development. Applying the production sequence-eligibility criteria yielded 9,613 proteins. Exact-sequence overlap with the evidence-supported reference and model-development populations was audited before inference, and structural annotations were retained for subsequent analysis.

All collections were processed using the frozen `phagelysclass_v1` bundle and ESM2-T30 embeddings generated with Sylphy using final-layer mean pooling and FP32 precision. Inference followed the frozen hierarchical architecture: global lytic recognition at a probability threshold of 0.5, followed by conditional multiclass assignment for globally positive proteins. Auxiliary subtype-compatibility scores were retained separately and did not modify the primary predictions. Source metadata were integrated only after inference. No application sequences were used for retraining, recalibration, threshold adjustment, or model selection. Consequently, application predictions were interpreted as source-specific classifier outputs, not as external estimates of predictive accuracy.

##### S1.7 Post-prediction physicochemical and geometric contextualization

Predictions from the three application collections were contextualized against the evidence-supported reference populations using a frozen panel of 26 sequence-derived descriptors. This analysis was independent of the ESM2-T30 representation used by the classifier and was performed only after the predictive architecture and application outputs had been frozen. Global lytic predictions were compared with the complete positive Core reference, whereas proteins receiving Endolysin, Depolymerase, or VAL assignments were compared with the corresponding clean single-class Core population.

For each reference population, the empirical distribution of every descriptor was used to define a 95% reference interval. Descriptor-level compatibility for application protein $i$ and descriptor $j$ was defined as

$$I_{ij} = \mathbb{I}\left(L_j \leq x_{ij} \leq U_j\right),$$

where $x_{ij}$ is the value of descriptor $j$ for protein $i$, and $L_j$ and $U_j$ are the lower and upper limits of the corresponding 95% reference interval. The fraction of descriptors compatible with the reference population was then calculated as

$$S_{\text{property}}(i) = \frac{1}{26}\sum_{j=1}^{26} I_{ij}.$$

A protein was considered to have property support when

$$S_{\text{property}}(i) \geq 0.80.$$

Descriptor-wise deviations from the reference distribution were additionally retained to identify which sequence properties contributed most strongly to disagreement. Population-level differences between application and reference proteins were summarized using the same effect-size framework used during the initial physicochemical characterization. These comparisons were descriptive and did not alter the predicted class.

Reference-space geometry was evaluated independently in the complete 26-dimensional descriptor space. Descriptor scaling was estimated from the corresponding evidence-supported reference population and subsequently applied without refitting to the application sequences. Euclidean distance was used throughout this standardized space. Quantitative geometric analyses were performed separately for the global positive reference and for the Endolysin, Depolymerase, and VAL reference populations.

For each application protein, geometric proximity was characterized using its distance to individual reference proteins, its local reference neighborhood, and its distance to the reference centroids. Empirical reference-distance distributions were used to define 95% and 99% coverage boundaries. The primary geometric-coverage criterion used the 95% boundary. For protein $i$ evaluated against reference population $c$,

$$S_{\mathrm{coverage}}(i, c) = I\left[d(i, c) \leq q_{0.95,c}\right],$$

where $d(i, c)$ denotes its reference-relative distance and $q_{0.95,c}$ is the corresponding empirical 95th-percentile reference boundary. The 99% boundary was retained as a secondary description of more extreme departures from reference space.

Subtype-specific local geometry was evaluated using three complementary assignments derived from the same standardized descriptor space: the majority subtype among the local reference neighbors, the subtype of the nearest individual reference protein, and the nearest subtype centroid. For a protein predicted as subtype $c$, local geometric support was assigned when at least two of these three criteria independently favored $c$,

$$S_{\mathrm{local}}(i, c) = I\left[I_{\mathrm{kNN}}(i, c) + I_{\mathrm{nearest}}(i, c) + I_{\mathrm{centroid}}(i, c) \geq 2\right].$$

This rule was used to distinguish membership within a broad subtype reference boundary from local agreement with the organization of the reference classes.

Low-dimensional projections were generated only to visualize these relationships. PCA, UMAP, and t-SNE were applied to reference and application proteins after descriptor-space standardization, but no distance, neighborhood, coverage, or support criterion was calculated from projected coordinates. All quantitative contextualization therefore remained defined in the complete 26-dimensional descriptor space.

The three post-prediction support axes were integrated at the individual-protein level. For a protein assigned to subtype $c$, the total support count was

$$C(i, c) = S_{\mathrm{property}}(i, c) + S_{\mathrm{coverage}}(i, c) + S_{\mathrm{local}}(i, c).$$

Proteins satisfying all three criteria ($C = 3$) were classified as *concordant*; proteins satisfying two criteria ($C = 2$) were classified as *mixed*; and proteins satisfying one or none ($C \leq 1$) were classified as *discordant*. These categories describe agreement between the frozen classifier, physicochemical properties, and reference-space geometry. They are not additional predictive classes and were not used to modify global or subtype predictions.

Exact sequence overlap between application proteins and the evidence-supported model-development reference sets was audited separately. Overlapping proteins were retained in the audit trail but excluded from concordance summaries intended to characterize behavior beyond exact reference-sequence reuse. The complete contextualization procedure was therefore applied only after prediction and remained independent of model training, architecture selection, threshold definition, and functional assignment.

##### S1.8 Implementation and reproducibility

PhageLysClass was implemented as a modular workflow with separate stages for data preparation, representation generation, model development, analysis, and deployment. Configuration files, execution scripts, intermediate results, and fitted models were preserved as independent artifacts to support reproducibility. The frozen production release, `phagelysclass_v1`, was exported separately from the benchmarking workflow. Its implementation specifications and software environment are summarized in Table **S12**.

Production provenance was recorded in `manifest.json`, including the frozen architecture, representation configuration, software environment, and SHA-256 references to the source artifacts and five task datasets used for final model fitting. File-level integrity was documented in `checksums.tsv` for all 35 release payloads. Each estimator underwent serialization round-trip validation, comparing predictions and probability

**Supplementary Table S12**: **Implementation and reproducibility specifications of the frozen PhageLysClass production release.**

| Element | Frozen specification |
|---|---|
| Model release | `phagelysclass_v1` |
| Architecture | `C_2_core_plus_3_aux` |
| Serialized estimators | 10 fitted estimators |
| Random state | 42 |
| Production representation | ESM2-T30, `facebook/esm2_t30_150M_UR50D` |
| Embedding contract | Final hidden layer, mean pooling, FP32, 640 dimensions |
| Sequence eligibility | 20 canonical amino acids, maximum length 1,024 residues |
| Model serialization | `joblib` estimators, JSON metadata, and ensemble recipes |
| Payload integrity | SHA-256 checksums for 35 payload files |
| Python | 3.12.14 |
| NumPy | 2.5.2 |
| pandas | 3.0.5 |
| scikit-learn | 1.9.0 |
| joblib | 1.5.3 |
| XGBoost | 3.4.1 |

estimates before and after reloading with a numerical tolerance of $10^{-10}$. Release integrity checks covered model artifacts, representation metadata, and checksums.

The standalone inference implementation supports CPU and CUDA execution and produces a machine-readable metadata sidecar documenting the model version, representation configuration, execution device, embedding batch size, sequence acceptance and rejection counts, and runtime. A positive-control FASTA was retained for inference smoke testing and was not used as an independent performance benchmark.

### S2 Supplementary Results

#### S2.1 Extended characterization of the evidence universe

The integration of PhageLysData and UniProtKB showed subtype-dependent differences in reference coverage and representation eligibility (Table **S13**). UniProtKB recovered a larger fraction of annotated Depolymerases than Endolysins or VALs. Representation eligibility was generally high across the evidence universe but decreased in the two challenge populations, primarily because of sequences exceeding the 1,024-residue limit.

**Supplementary Table S13**: **Reference coverage and representation eligibility across the PhageLysClass evidence universe.**

| Population | Sequences ($n/N$) | Fraction (%) |
|---|---|---|
| *Core overlap with UniProtKB* | | |
| Endolysin | 5,294 / 9,888 | 53.5 |
| Depolymerase | 540 / 712 | 75.8 |
| VAL | 125 / 241 | 51.9 |
| *ESM2 representation eligibility* | | |
| Positive Core | – | 94.9 |
| Easy negatives | – | 98.5 |
| Hard negatives | – | 99.9 |
| Explicit unlabeled | – | 99.6 |
| Other unresolved | – | 97.0 |
| Lytic-related challenge | 270 / 319 | 84.6 |
| Ambiguous structural challenge | 9,613 / 10,885 | 88.3 |

Representation-space analysis further revealed differences in the geometric separation of Core positives from the supervised non-target backgrounds. In the alternative-function comparison, silhouette scores were 0.048 for sequence composition, 0.082 for physicochemical descriptors, and 0.058 for their combined representation. Separation increased with ESM2-T6 (0.101) and ESM2-T30 (0.144). A similar pattern was observed against the mixed background, with silhouette scores of 0.097 and 0.139 for ESM2-T6 and ESM2-T30, respectively.

These relatively low values indicate substantial overlap between positive and non-target populations, even in protein language model representation spaces.

The lysis-adjacent background showed consistently greater separation from Core positives across the evaluated representations. With ESM2-T30, its silhouette score reached 0.312, compared with 0.144 for the alternative-function background and 0.139 for the mixed background. Thus, the biologically closer lysis-adjacent population was not necessarily the most geometrically similar to Core positives. These findings highlight differences between biological task definitions and the organization of proteins in numerical representation spaces.

#### S2.2 Complete factorial analysis of predictive generalization

The factorial analysis examined task-specific performance distributions, changes in available sequence evidence, and the sensitivity of generalization estimates to redundancy and train–test homology control. The complementary results are summarized in Tables **S14**–**S16**.

##### S2.2.1 Performance distributions across tasks and sequence-space conditions

Predictive performance differed across tasks and sequence-space conditions (Table **S14**). Global hard recognition retained the highest median MCC under both R90/S90 (0.940) and R30/S30 (0.874), whereas Endolysin and VAL showed greater deterioration under stricter conditions. Depolymerase remained comparatively stable, although its small positive redundancy contrast was not preserved in the common-test analysis. VAL exhibited the greatest performance dispersion, consistent with its limited positive reference population.

**Supplementary Table S14**: **Supervised predictive performance across the factorial generalization benchmark.**

| Task | R90/S90 MCC | R30/S30 MCC | $\Delta R$ | $\Delta S$ | $\Delta$ joint |
|---|---|---|---|---|---|
| Global easy | 0.775 [0.697, 0.912] | 0.665 [0.460, 0.809] | −0.102 | −0.018 | −0.131 |
| Global hard | 0.940 [0.872, 0.957] | 0.874 [0.796, 0.901] | −0.046 | −0.007 | −0.064 |
| Global mixed | 0.783 [0.692, 0.900] | 0.649 [0.453, 0.788] | −0.103 | −0.023 | −0.127 |
| Endolysin | 0.775 [0.688, 0.914] | 0.590 [0.377, 0.758] | −0.143 | −0.019 | −0.174 |
| Depolymerase | 0.869 [0.746, 0.911] | 0.849 [0.688, 0.913] | +0.018 | −0.008 | +0.000 |
| VAL | 0.714 [0.426, 0.833] | 0.406 [0.110, 0.675] | −0.044 | −0.035 | −0.194 |
| Multiclass clean | 0.841 [0.751, 0.887] | 0.821 [0.681, 0.859] | −0.023 | −0.020 | −0.030 |

Across the 140 task–representation–learner surfaces, the median joint R90/S90-to-R30/S30 change was −0.084 MCC. However, the task-specific contrasts showed that the generalization penalty was not uniform, particularly for Endolysin, Depolymerase, and VAL.

##### S2.2.2 Redundancy reduction and evidence retention

Redundancy and homology control affected different properties of the evaluation datasets. Table **S15** summarizes changes in sequence retention and homology-cluster structure across 112 unique supervised task–R–S dataset states, independently of the predictive strategies evaluated within each state.

**Supplementary Table S15**: **Changes in evidence quantity and sequence-space structure under redundancy and homology control.**

| *Redundancy reduction relative to R90* | | | | |
|---|---|---|---|---|
| **Regime** | **Train entities** | **Positive support** | **Cluster richness** | **Entities/cluster** |
| R90 | 1.000 | 1.000 | 1.000 | 1.000 |
| R70 | 0.717 | 0.737 | 1.000 | 0.721 |
| R50 | 0.497 | 0.508 | 0.856 | 0.624 |
| R30 | 0.307 | 0.337 | 0.560 | 0.624 |
| *Homology separation relative to S90* | | | | |
| **Regime** | **Train entities** | **Test entities** | **Train clusters** | **Entities/cluster** |
| S90 | 1.000 | 1.000 | 1.000 | 1.000 |
| S70 | 1.000 | 1.000 | 0.998 | 1.001 |
| S50 | 1.000 | 1.000 | 0.858 | 1.195 |
| S30 | 1.000 | 0.999 | 0.554 | 1.824 |

At R30, the median retained fractions of training entities and positive sequences were 30.7% and 33.7%, respectively, whereas relative cluster richness remained at 56.0%. Redundancy reduction therefore removed sequence entities more rapidly than homology groups, although increasingly strict reduction also decreased the number of represented clusters. In contrast, S90-to-S30 homology separation largely preserved dataset size while reducing the number of training clusters and increasing their average membership. These differences explain why redundancy reduction and homology-aware partitioning cannot be treated as equivalent sequence-control operations.

###### S2.2.3 Homology effects, common-test sensitivity, and redundancy–homology interaction

Native-test comparisons indicated a stronger performance penalty from redundancy reduction than from homology separation. Because independently generated R–S conditions could contain different test proteins, common-test comparisons were used to determine whether these effects persisted when models were evaluated on shared sequences (Table **S16**).

**Supplementary Table S16**: **Native-test and common-test sensitivity of extreme redundancy and homology contrasts, together with task-specific R–S interactions.**

| Task | $\Delta R_{\text{native}}$ | $\Delta R_{\text{common}}$ | $n_R$ | Min. support | $\Delta S_{\text{native}}$ | $\Delta S_{\text{common}}$ | Interaction | Positive (%) |
|---|---|---|---|---|---|---|---|---|
| Global easy | −0.102 | −0.022 | 80 | 143.0 | −0.018 | −0.019 | +0.047 | 95 |
| Global hard | −0.046 | −0.012 | 80 | 64.5 | −0.007 | −0.000 | +0.021 | 80 |
| Global mixed | −0.103 | −0.016 | 80 | 136.5 | −0.023 | −0.015 | +0.010 | 70 |
| Endolysin | −0.143 | −0.032 | 80 | 110.0 | −0.019 | −0.011 | +0.080 | 95 |
| Depolymerase | +0.018 | −0.006 | 80 | 28.0 | −0.008 | −0.014 | +0.061 | 90 |
| VAL | −0.044 | +0.000 | 60 | 6.5 | −0.035 | +0.000 | +0.339 | 100 |
| Multiclass clean | −0.023 | +0.000 | 80 | 5.5 | −0.020 | −0.014 | +0.029 | 90 |

Across the 540 calculable extreme redundancy contrasts, the median native-test MCC difference was approximately −0.075, compared with −0.012 on shared test proteins. Endolysin retained the largest common-test penalty (−0.032), while the apparent native-test improvement for Depolymerase (+0.018) became slightly negative (−0.006). These results indicate that changes in test-set composition contributed substantially to the observed redundancy penalty, although performance differences persisted for some tasks.

The homology-separation effect was less sensitive to test composition but depended on the existing redundancy regime. At R90, the native and common-test S30-minus-S90 contrasts were approximately −0.047 and −0.037, respectively, whereas both approached zero at R30. Paired bootstrap analysis supported this attenuation, with the fraction of common-test S contrasts showing confidence intervals entirely below zero decreasing from approximately 48% at R90 to 2% at R30. These intervals describe uncertainty conditional on the fitted models and shared evaluation sequences.

The factorial interactions were predominantly positive. Across 140 supervised surfaces, the median extreme R–S interaction was +0.051 MCC, with 88.6% of surfaces showing a positive interaction. The corresponding task-level estimates are provided in Table **S16**. An independent decomposition attributed median fractions of 70.5%, 9.1%, and 18.3% of within-surface variation to redundancy, homology separation, and their interaction, respectively. Thus, redundancy was the dominant factor, but its interaction with homology separation accounted for a larger fraction of variation than the independent S effect.

These findings require consideration of common-test support. Extreme R and S intersections retained approximately 28% and 30% of the smaller original test populations, respectively. For VAL, only 60 of 80 extreme R contrasts yielded calculable common-test MCC, with a median minimum class support of 6.5 proteins. Moreover, four-corner intersections retained a median of only 74 proteins, preventing reliable estimation of the complete factorial interaction on identical test sequences. Consequently, the interaction estimates characterize the native benchmark surfaces, whereas the common-test analyses provide more direct evidence about prediction differences on shared proteins.

##### S2.3 Representation, learner, and positive–unlabeled behavior

The factorial benchmark was further analyzed to quantify representation-dependent robustness, learner ranking stability, and the effects of positive–unlabeled learning on unresolved-sequence prioritization.

###### S2.3.1 Representation-dependent performance and robustness

The representation-level robustness analysis is summarized in Table **S17**. ESM2-T30 ranked first in 92.9% of matched comparisons and showed the highest worst-case MCC (0.854), together with the smallest median sensitivity to redundancy ($|\Delta R| = 0.062$) and homology separation ($|\Delta S| = 0.015$). Its median MCC range across the complete R–S surfaces was 0.093, compared with 0.118 for ESM2-T6. These results indicate that the representation advantage extended beyond average predictive performance to stability across evaluation conditions.

**Supplementary Table S17: Predictive performance and robustness by protein representation.**

| Representation | Median MCC | Worst MCC | R30/S30 MCC | First (%) | Median $\lvert\Delta R\rvert$ | Median $\lvert\Delta S\rvert$ |
|---|---|---|---|---|---|---|
| One-hot | 0.496 | 0.380 | 0.390 | 0.0 | 0.099 | 0.038 |
| Roxy | 0.675 | 0.569 | 0.616 | 0.0 | 0.087 | 0.036 |
| ESM2-T6 | 0.829 | 0.758 | 0.782 | 6.6 | 0.083 | 0.020 |
| ESM2-T30 | 0.902 | 0.854 | 0.881 | 92.9 | 0.062 | 0.015 |

Paired bootstrap comparisons further characterized the statistical support for these differences. ESM2-T30 improved MCC over Roxy by a median of 0.214, with 95% confidence intervals entirely above zero in 95% of 140 evaluated contrasts. The T30–T6 difference was smaller (+0.069 MCC), with 59% of intervals entirely above zero. Support for the latter advantage was task-dependent, particularly for global easy, global mixed, and Endolysin recognition, whereas most global-hard comparisons were statistically unresolved.

##### S2.3.2 Learner effects and ranking stability

The supervised learner comparison is summarized in Table **S18**. MLP, XGBoost, and KNN formed a competitive group, with median gaps to the best learner of 0.008, 0.012, and 0.014 MCC, respectively. The median MLP–XGBoost difference was only +0.006 MCC, and approximately 89% of their paired bootstrap confidence intervals overlapped zero. Logistic regression exhibited the smallest median sensitivity to R and S but lower overall and worst-case performance, illustrating that limited performance variation did not necessarily imply stronger predictive generalization.

**Supplementary Table S18: Supervised learner performance and robustness across representations and R–S conditions.**

| Learner | Median MCC | Worst MCC | First (%) | Gap to best | Median $\lvert\Delta R\rvert$ | Median $\lvert\Delta S\rvert$ |
|---|---|---|---|---|---|---|
| LR | 0.717 | 0.665 | 11.8 | 0.075 | 0.063 | 0.018 |
| RF | 0.742 | 0.677 | 4.9 | 0.064 | 0.091 | 0.022 |
| KNN | 0.784 | 0.701 | 27.2 | 0.014 | 0.088 | 0.032 |
| XGBoost | 0.796 | 0.718 | 21.0 | 0.012 | 0.079 | 0.023 |
| MLP | 0.817 | 0.752 | 31.9 | 0.008 | 0.076 | 0.022 |

Learner dependence decreased with representation quality. The median MCC range between the best and worst learner declined from 0.306 with one-hot encoding to 0.051 with ESM2-T30. Across the 112 task–R–S conditions, the representation-dependent performance range exceeded the learner-dependent range in every condition, with a median ratio of approximately 2.8. Furthermore, the highest-performing learner changed between R90/S90 and R30/S30 in 18 of the 28 task–representation surfaces (64.3%). These findings support endpoint-specific learner selection based on the complete generalization landscape.

##### S2.3.3 Positive–unlabeled learning and unresolved-sequence prioritization

The PU benchmark comprised 768 experiments across two unlabeled universes, four representations, two base learners, three learning paradigms, and 16 R–S conditions. Evaluation focused on recovery of known positives and ranking of unresolved proteins. U95 recovery was defined as the fraction of held-out known positives scoring at or above the 95th percentile of the corresponding unlabeled-score distribution. Because unlabeled proteins were not confirmed negatives, this measure was interpreted as a ranking diagnostic rather than biological sensitivity.

Known-positive recovery depended primarily on representation quality (Table **S19**). ESM2-T30 achieved median U95 recovery values near 0.93 across all three PU paradigms, with little advantage from BaggingPU or Elkan–Noto correction. BaggingPU produced its largest recovery improvement under one-hot encoding (+0.093), while its median paired gain was negligible or slightly negative for the remaining representations.

Similar positive recovery did not imply equivalent candidate rankings. With ESM2-T30, naive P-versus-U and BaggingPU produced nearly identical U95 recovery, but their unlabeled rankings showed a median Spearman correlation of 0.808 and a top-5% Jaccard overlap of 0.530. Elkan–Noto correction, in contrast,

**Supplementary Table S19**: **Known-positive recovery and unlabeled-ranking changes across PU learning paradigms.**

| Representation | Naive U95 | Bagging U95 | EN U95 | Bagging gain | U Spearman | Top-5% Jaccard |
|---|---|---|---|---|---|---|
| One-hot | 0.454 | 0.567 | 0.493 | +0.093 | 0.669 | 0.410 |
| Roxy | 0.709 | 0.711 | 0.709 | +0.000 | 0.968 | 0.812 |
| ESM2-T6 | 0.852 | 0.849 | 0.852 | −0.003 | 0.908 | 0.701 |
| ESM2-T30 | 0.930 | 0.931 | 0.930 | +0.000 | 0.808 | 0.530 |

largely preserved the naive ranking while modifying score scaling. Its corrected upper-tail thresholds frequently reached 1.0 because of probability clipping, limiting the interpretation of apparent improvements in extreme positive recovery.

PU recovery also remained sensitive to sequence-space control. The median R90-to-R30 change in U95 recovery was approximately −0.073, compared with −0.008 for S90-to-S30, with approximately 90% of PU performance surfaces classified as R-dominant. The modest relative benefit of BaggingPU at R30 did not compensate for the overall reduction in known-positive recovery.

Expanding the unlabeled universe from `U_explicit` to include `U_other_unresolved` introduced additional differences in recovery and candidate selection (Table **S20**). Native-test U95 recovery decreased under all three paradigms, but the differences were substantially attenuated when evaluation was restricted to shared positives and the original `U_explicit` reference population.

**Supplementary Table S20**: **Effect of expanding the PU unlabeled universe on known-positive recovery and candidate prioritization.**

| PU method | Native ΔU95 | Common-reference ΔU95 | Other-U in top 5% | Enrichment | Top-1% Jaccard |
|---|---|---|---|---|---|
| Naive P-versus-U | −0.054 | −0.009 | 53% | 1.90 | 0.46 |
| BaggingPU | −0.056 | ≈ 0.000 | 58% | 2.07 | 0.53 |
| Elkan–Noto | −0.047 | −0.009 | 53% | 1.90 | 0.48 |

The additional unresolved population contributed 53–58% of the highest-scoring 5% of broad-U proteins, corresponding to approximately 1.9–2.1-fold enrichment relative to its prevalence in the evaluated unlabeled universe. Nevertheless, expanding the unlabeled population also altered the ranking of proteins already present in `U_explicit`. Conservative–broad top-1% Jaccard overlap ranged from approximately 0.46 to 0.53, indicating substantial differences in candidate selection even when evaluating a shared sequence population.

Overall, the PU analyses distinguished recovery of known positives from prioritization of unresolved sequences. Representation quality primarily influenced the former, while learning paradigm and unlabeled-universe definition influenced the latter. These differences characterize alternative prioritization regimes without establishing biological function or calibrated probabilities of lytic activity.

##### S2.4 Predictive-strategy selection and architecture construction

Predictive strategies were selected independently for each supervised endpoint using their performance and robustness across the complete factorial benchmark. A strategy was retained if it led at least one predefined operational criterion or achieved a mean MCC within 0.02 of the best strategy for its task. Selection criteria included mean, worst-case, and strict R30/S30 performance, rank consistency, regret, sensitivity to sequence-space restrictions, and computational efficiency. No aggregate selection score was constructed. This procedure retained 23 individual strategies, comprising 20 ESM2-T30 and three ESM2-T6 models (Table **S21**).

All non-singleton combinations within the retained portfolios were evaluated, yielding 46 ensemble candidates and 736 R–S evaluations for each of the unweighted and validation-weighted soft-voting formulations. Ensemble gains were generally modest and endpoint-dependent (Table **S22**). The selected global and Endolysin ensembles improved mean, worst-case, and strict-condition MCC relative to their corresponding individual baselines. The multiclass XGB–LR ensemble provided the largest strict-condition improvement (+0.0197 MCC), whereas the selected Depolymerase ensemble improved strict performance without improving the best individual worst-case result. Comparisons were criterion-specific, and the strongest individual baseline could differ between metrics.

**Supplementary Table S21**: **Individual predictive strategies retained after factorial robustness analysis.**

| Endpoint | Candidates | Retained strategies | Production role |
|---|---|---|---|
| Global easy | 3 | T30–KNN; T30–XGB; T30–MLP | Evaluation only |
| Global hard | 4 | T30–RF; T6–KNN; T30–XGB; T30–MLP | Evaluation only |
| Global mixed | 3 | T30–KNN; T30–MLP; T30–XGB | Global recognition core |
| Endolysin | 3 | T30–KNN; T30–XGB; T30–MLP | Auxiliary compatibility |
| Depolymerase | 4 | T30–MLP; T30–KNN; T6–KNN; T6–MLP | Auxiliary compatibility |
| VAL | 2 | T30–MLP; T30–LR | Auxiliary compatibility |
| Multiclass clean | 4 | T30–MLP; T30–XGB; T30–KNN; T30–LR | Primary functional core |
| **Total** | **23** | **20 T30 strategies and 3 T6 strategies** | |

**Supplementary Table S22**: **Selected predictive recipes and their performance differences relative to individual models.**

| Component | Selected recipe | Mean MCC | Δ mean | Δ worst | Δ strict |
|---|---|---|---|---|---|
| Global core | T30–KNN + T30–MLP | 0.9280 | +0.0045 | +0.0104 | +0.0052 |
| Multiclass core | T30–XGB + T30–LR | 0.8899 | +0.0028 | +0.0041 | +0.0197 |
| Endolysin auxiliary | T30–KNN + T30–XGB + T30–MLP | 0.9288 | +0.0066 | +0.0034 | +0.0048 |
| Depolymerase auxiliary | T30–MLP + T30–KNN | 0.9241 | +0.0042 | −0.0002 | +0.0105 |
| VAL auxiliary | T30–MLP | 0.8578 | – | – | – |

Ensembling was not uniformly beneficial. For VAL, the unweighted MLP–LR ensemble increased mean MCC by approximately 0.0053 but reduced strict-condition MCC by 0.0416 relative to the strongest individual model at R30/S30. VAL therefore retained a single MLP classifier. Validation-weighted voting provided no material advantage over uniform averaging, supporting the adoption of unweighted soft voting for the remaining components. Prediction disagreement alone was also a poor indicator of ensemble improvement (Spearman $\rho \approx 0.06$).

Three architectural organizations were subsequently evaluated against eight predefined constraints concerning prediction hierarchy, subtype compatibility, output precedence, and inference simplicity. The two-core organization satisfied seven constraints but omitted independent subtype-compatibility outputs, whereas the five-module co-equal architecture satisfied only three because of conflicting prediction roles. The two-core-plus-three-auxiliary organization satisfied all eight constraints and was selected as the production architecture.

The selected recipes were subsequently consolidated into the frozen production configuration (Table **S23**). For each retained learner, the modal configuration selected through inner validation across the 16 R–S conditions was adopted, with validation performance used to resolve ties. Final estimators were refitted using all eligible sequences from their corresponding task definitions, without retaining an outer benchmark test partition. No application sequence contributed to configuration selection or refitting.

**Supplementary Table S23**: **Frozen PhageLysClass production components and refitting configurations.**

| Component | Tier | Members and frozen configurations | Fit proteins | Estimators |
|---|---|---|---|---|
| Global lytic recognition | Core | KNN02 + MLP02 | 69,448 | 2 |
| Clean multiclass function | Core | XGB01 + LR01 | 10,244 | 2 |
| Endolysin compatibility | Auxiliary | KNN01 + XGB01 + MLP02 | 67,560 | 3 |
| Depolymerase compatibility | Auxiliary | MLP01 + KNN03 | 58,824 | 2 |
| VAL compatibility | Auxiliary | MLP01 | 58,427 | 1 |
| **Total fitted estimators** | | | | **10** |

The strict R30/S30 performance results presented in the main text correspond to the selected benchmark recipes before full-data production refitting. These measurements contributed to architecture selection and therefore do not constitute independent post-selection validation. This distinction is particularly relevant for VAL, which had only 17 positive proteins in its strict binary test set, while the smallest class in the clean multiclass test contained 14 proteins. Dedicated post-selection confidence intervals for the refitted production ensembles were not established.
